# Addressing reverse inference in structural brain alterations

**DOI:** 10.1101/536847

**Authors:** Franco Cauda, Andrea Nani, Donato Liloia, Jordi Manuello, Enrico Premi, Sergio Duca, Peter T. Fox, Tommaso Costa

**Affiliations:** GCS- fMRI, Koelliker Hospital and Department of Psychology, University of Turin, Turin, Italy; Department of Psychology, University of Turin, Turin, Italy; FOCUS Lab, Department of Psychology, university of Turin, Turin, Italy; Stroke Unit, Azienda Socio Sanitaria Territoriale Spedali Civili, Spedali Civili Hospital, Brescia, Italy; Centre for Neurodegenerative Disorders, Neurology Unit, Department of Clinical and Experimental Sciences, University of Brescia, Brescia, Italy; Research Imaging Institute, University of Texas Health Science Center at San Antonio, USA; South Texas Veterans Health Care System, San Antonio, Texas, USA

**Keywords:** Brain disorders, Voxel-based morphometry, Alteration specificity, Bayes’, factor, Reverse probability, Schizophrenia, Alzheimer’s disease, Pain

## Abstract

In the field of neuroimaging reverse inferences can lead us to suppose the involvement of cognitive processes from certain patterns of brain activity. Still, the same reasoning holds if we substitute “brain activity” with “brain alteration” and “cognitive process” with “brain disorder”. To assess the involvement of a certain alteration pattern in a brain disorder we used the Bayes’ factor technique on voxel-based morphometry data of schizophrenia and Alzheimer’s disease. This technique allows to calculate the ratio between the likelihoods of two alternative hypotheses (in our case, that the alteration of the voxel is specific for the brain disorder under scrutiny or that the alteration is not specific). We then performed temporal simulations of the alterations’ spread associated with different pathologies. The Bayes’ factor values calculated on these simulated data were able to reveal that the areas, which are more specific to a certain disease, are also the ones to be early altered.

## Introduction

The study of the cerebral areas that are co-altered in the pathological brain is of fundamental importance to better understand how neuropathologies spread and develop. This knowledge is essential especially with regard to mental illnesses, as it could significantly improve their classification as well as their diagnoses (1). The psychopathological models DSM-based (2) and the corresponding classification of the World Health Organization (3) consider psychiatric and neurological conditions as distinct clinical constructs with different etiologies and biomarkers. Still, this view, which is mainly derived from clinical observations as well as from patient self-reports, may be partially mistaken, as traditional categorizations might regard as similar certain clinical manifestations that are, on the contrary, characterized by biological heterogeneity, and classify as different certain disorders that, instead, may share neurobiological substrates.

The traditional classification approach has led to the idea that each brain disease is real, well defined and caused by specific factors. However, growing evidence is starting to challenge this view (4, 5). Large-scale phenotypic studies, for example, provide evidence that brain disorders share different liabilities (4, 6). Moreover, etiological investigations give further support for ruling out any rigid classification of mental illnesses, which are frequently characterized by polygenic inheritance with multiple small-effect risk alleles causing a constant diffusion of genetic liability (7). Comorbidity, too, defies a rigid categorization. Co-occurrences of psychiatric diseases, in fact, are rather frequent than exceptional (8, 9). Therefore, this large diversity in symptomatology, dimensionality and comorbidity (10-12) suggests that our models of classification need to be profoundly revised (13).

A transdiagnostic approach to brain disorders is supported by several studies showing that a common set of brain regions is generally altered in a variety of psychiatric (4, 14-18) as well as neurological (8) conditions. As shown in Cauda et al. (8), most of brain disorders are likely to produce anatomical alterations that largely overlap with each other. This study has analyzed the voxel-based morphometry (VBM) database of BrainMap and created an alteration map showing the alteration rate of brain disorders, that is, the percentage of how many pathologies affect every area of the brain (Fig. 1). BrainMap is an online open access neuroimaging data repository that is based on a systematic coding scheme; it stores thousands of published human neuroimaging experimental results, as well as reports thousands brain locations in stereotaxic space. The VBM dataset of BrainMap is therefore an ideal environment for carrying out structural meta-analyses (19-22).

The maps of the top row of Fig. 1 reports in percentage how many pathologies of the VBM BrainMap database cause alterations in every area of the brain: this map demonstrates that there is a common set of altered regions (such as the insulae, the anterior cingulate cortex, some of the prefrontal and anterior temporal areas) which is affected by the majority of brain disorders (in these areas more than 90% of pathologies have at least one focus of alteration). This large overlap of areas altered by different brain disorders makes them scarcely informative about the development of neuropathological processes. In other words, in these areas the alteration pattern is rather non-specific. Moreover, our study (8) shows that there are almost no brain areas specifically altered by one pathology only. There are some areas, however, that are altered by a limited number of diseases and can therefore be considered as more specific to certain brain disorders (see (8) and Fig. 1 of this study). This fact makes very less informative the analyses based on “forward” inferences (see below) (23), as they tend to treat in the same way both the areas having high or low specificity. This issue can however be solved, at least partially, through a Bayesian reverse inference approach.

The temporal progression of neuropathology, which develops from a limited number of alterations to a larger number of altered areas (24), implies that, most likely, reverse inferences identify as more specific to a certain brain disorder the earliest areas to be affected. Theoretically, the maximum level of overlap between altered regions reaches its limit when all the brain is affected. In this hypothetical case, each pathology would alter most of the cerebral areas. In fact, as alterations gradually spread, the overlap of alterations caused by different pathologies will be greater and greater, thus reducing the degree of specificity of the areas that are progressively more altered (for an infographic see Fig. S4 of the Supplementary online Materials). Nevertheless, also the non-linear progression of brain alterations (both in pathological and in healthy ageing individuals) as well as the largely unknown complexity (apart from age and gender) of potential modulating factors (i.e., APOE and other genetic and environmental factors, etc.) should be taken into account (25-27).

Given the premises discussed so far, our study aims to address four important points for the understanding of the pathological brain as well as for the improvement of the approach to the technique of the reverse inference. First, we put forward a map based on a Bayesian reverse inference of the two most represented brain diseases in BrainMap (schizophrenia (SCZ) and Alzheimer’s disease (AD)). Second, we propose solutions to some critical points about the Bayesian reverse inference approach. Third, we discuss how to tackle the potential bias of the inhomogeneous representativeness of pathologies within the dataset of BrainMap. Fourth, we investigate how much the inhomogeneity of the sample may influence the calculus of the Bayesian reverse inference. Fifth, we put forward a simulation of a series of pathologies and the spread of their alterations so as to see how a calculus based on a Bayesian reverse inference can identify which brain areas are the earliest to be affected. Let us see now each of these points in more details.

1) To create, through a Bayesian reverse inference approach, a map of the two most represented pathologies in BrainMap (i.e., SCZ and AD) and identify the areas, if any, that are more specific for these two disorders using as input data the maps obtained with the activation likelihood estimation (ALE) method.

Researchers generally apply a forward inference by asking what are the areas affected by a certain pathology; in our case, we are instead interested in the reverse inference by asking what are the pathological conditions that might have produced a specific alteration pattern. In fact, the probability that a specific pattern of gray matter (GM) alterations be related to a specific brain disorder is not equivalent to the probability that a specific brain disorder be related to a specific pattern of GM alterations: P(GM alteration|brain disease) ≠ P(brain disease|GM alteration). To our best knowledge, this kind of calculus has never been tried before on meta-analytical data of morphological alterations but only on healthy subjects’ task-based fMRI data (23, 28, 29). We know about a study that has tried a type of reverse inference over functional data derived from tasks performed by groups of patients suffering from psychiatric conditions (30). However, this study is not based on a Bayesian reverse inference: authors applied a Chi squared test with the Yates’ correction on a cross tables diagnosis-by-region and tested whether studies on a pathology report similar results to another one. In sum, the authors calculated a form of correlation between the number of studies of the pathology-by-region to those of another pathology.

2) To contribute to the development of the method of reverse inference by trying to address some of the critical issues of this approach.

The possibility of the reverse inference from neuroimaging data has already been explored (23, 29, 31-34). On the legitimacy of the method *per se* see Lieberman and Eisenberger (35), Poldrack (36), Lieberman (37), Yarkoni (38), Yarkoni (39), ShackmanLab (40), Wager, Atlas (41), Gelman (42), Machery (31) and Hutzler (32). With a pioneering study, Poldrack (23) has highlighted the difficulties of this kind of inference in the field of functional neuroimaging. Very specific patterns of activations associated with pathological conditions are extremely infrequent (29, 43). Difficulties in making a reverse inference correctly have been extensively discussed in the debate raised after the publication of Lieberman and Eisenberger (35) (36-42), who claimed that activity of the anterior dorsal cingulate is selective for pain. Results put forward by these authors are based on analyses performed with the help of Neurosynth (28), a database containing coordinates of activation and terms of different cognitive domains derived from a large set of studies (11000). These findings have been criticized by Yarkoni, and other authors (36-42).

The theoretical framework of the reverse inference is the Bayes’ theorem. More specifically, through a reverse inference we can infer the posterior probability of a certain cognitive process M starting from a pattern of brain activation A. Thus, this inference is based on the conditional probability or likelihood P(A|M) and a prior probability p(M), that is, the probability that we have before acquiring any clues on brain activations. Neuroimaging data can therefore provide information on the likelihood of M given A. And we can go through the same reasoning if we substitute “cognitive process M” with “brain pathology M” and “brain activation A” with “brain alteration A”. In fact, VBM data can provide information on the likelihood of a brain pathology given a specific pattern of GM alterations.

Poldrack (23) used the BrainMap database (19) to estimate the activation of Broca’s area in supporting language as an example of reverse inference. More specifically, he calculated the number of studies in which Broca’s area was active when involved in linguistic tasks and the number of studies in which Broca’s area was active when involved in non-linguistic tasks. Then, on the basis of the prior probability of the “cognitive process M” *P*(*M*) = 0.5 (i.e., the equal probability to be activated or not activated by the cognitive process), he found a posterior probability of 0.69 that language is supported by Broca’s area activation. To better assess the involvement of a certain brain activation in a cognitive process we can use the Bayes’ factor (BF) (44), which is the ratio between the likelihoods of two alternative hypotheses (see the Materials and methods section for an in-depth description of this index). In Poldrack’s case, the BF is 2.3, which is considered to be a weak evidence of specificity (44). Poldrack (29) has also claimed that the BrainMap database may be biased by the fact that the studies are introduced manually, as this could lead to a partial sample of the literature. In contrast, the database of Neurosynth, which uses an automated process for selecting articles, should have a more comprehensive sample. We agree only in part with this criticism. In fact, even if we could create an all-inclusive database of task-based fMRI studies, the bias due to the fact that the various cognitive domains are not uniformly represented would not be definitely ruled out. This is because all the topics of research do not have the same interest for the scientific community: for instance, i) some tasks or pathologies are more or less studied than others, independently of their relative frequency; ii) some other task or pathologies are more difficult to study and, therefore, less investigated; iii) the automated selection process for introducing experiments in a database can make classification mistakes, which are sometimes more frequent than those occurring with a non-automated selection; iv) Neurosynth does not differentiate between activation and deactivation. In any case, the aim of the present study is not to explore all the strengths and weaknesses of databases, such as BrainMap or Neurosynth, but rather to propose solutions and, at least in part, try to overcome the problems raised in the debate about the reverse inference.

The choice of prior probabilities is probably the most complex issue in the Bayesian approach to meta-analytical data. Priors are generally set to 0.5, so that neither hypothesis *P*(*H*_1_) nor hypothesis *P*(*H*_2_) is privileged, where *H*_1_is the hypothesis of specificity and *H*_2_ the hypothesis of non-specificity, respectively. However, Yarkoni has claimed in a series of commentaries (28, 39) that it would be more convenient to choose different priors, given that the base rates related to pathologies are different. As we propose to show in this study (see the Materials and methods section), it is our advice that this suggestion is correct for the priors regarding the parameters of the likelihood model rather than for the priors regarding the hypotheses. In fact, in all the Bayesian and frequentist statistics, the likelihood is dependent on the number of the samples. In contrast, the priors regarding the hypotheses are independent of the numerosity of the samples, as an impartial position regarding the two hypotheses is usually assumed; this is true for both Bayesian and frequentist statistics. Moreover, it should be noted that with a sufficient amount of data, as it is our case, the BF should converge, at least in theory, to the value “true”, even though the priors are supposed to be equiprobable. It is in fact plausible that, even though you and I can have different prior beliefs, more often we will agree over the form of the likelihood, so that, if we gather enough data, the posterior will become very close (45). In any case, to avoid the introduction of prior densities regarding the parameters, and to assess whether or not the use of inhomogeneous data may produce incorrect BF values, we propose to apply the Schwarz criterion (46, 47), which, for large samples, can be considered as an approximation of the logarithm of BF. In fact, as we will see in the Methods section, the Schwarz criterion maintain a strict relationship with the Bayesian information criterion (BIC), which depends on the numerosity of the sample, so that we can control through it how much the BIC is close to the BF with regard to the numerosity of the sample. To do so we calculated the BIC (see the Materials and methods section) on functional data (i.e., pain condition) and compared these results with those obtained with the BF determined with equiprobable priors. As postulated by the Schwarz criterion, our hypothesis is that the BF values, calculated with equiprobable priors, converge to the BIC if they do not suffer from the bias due to the inhomogeneity of the sample affecting the priors of the parameters of the likelihood model.

3) To reduce the potential bias of the inhomogeneous representativeness of pathologies within the BrainMap database.

This strategy is directed to minimize the problem of inhomogeneity by recalculating the BF values with an ALE that include all the data related to each pathology of the database, so as to have only one sample for each pathology to be compared with the original result. To do so we created a single sample for every pathology by constructing an ALE map of all the experiments about each specific pathology. The “compensated” BF maps so obtained were then correlated to other maps constructed without this compensation in order to understand how much the inhomogeneous representativeness of pathologies can influence the BF maps.

4) To inspect how much the inhomogeneity of the sample, which can be due to the manual selection of the data as well as to the uneven representation of brain pathologies in the database, can influence the BF calculus.

This was performed by testing our results by means of injection of different amount of spurious data in the analysis.

5) To find out whether the BF calculus can detect which cerebral areas are the earliest to be altered.

This was achieved by means of a simulation of a series of pathologies and of their different stages of alteration spread.

## Materials and methods

### Selection of studies

A multiple systematic search was conducted in the BrainMap database (http://brainmap.org/) (19, 21, 22) to identify neuroimaging experimental results of interest. BrainMap is an online open access database of published functional and VBM experiments that reports both the coordinate-based results (x,y,z) in stereotactic brain space and a hierarchical coding scheme of meta-data concerning the experimental methods and conditions (20). At the time of the selection phase (April 2018), BrainMap included meta-data associated with more than 4000 publications, containing over 19000 experiments, 148000 subjects and 149000 coordinate-based results.

First, we queried the VBM BrainMap database sector (22) in order to identify the two of the most represented brain disorders of this structural dataset (i.e., SCZ and AD). By means of the software application Sleuth 2.4, we employed a double systematic search to retrieve the eligible voxel-based results for each of the two brain disorders. The search algorithms were constructed as follows:

For the meta-analysis of SCZ:

SCZ QUERY A) [*Experiments Context IS Disease] AND [Experiment Contrast IS Gray Matter] AND [Experiments Observed Changes IS Controls>Patients] AND [Experiments Observed Changes IS Controls<Patients] AND [Subjects Diagnosis IS Schizophrenia]*;

SCZ QUERY B) [*Experiments Context IS Disease] AND [Experiment Contrast IS Gray Matter] AND [Experiments Observed Changes IS Controls>Patients] AND [Experiments Observed Changes IS Controls<Patients] AND [Subjects Diagnosis IS NOT Schizophrenia]*.

For the meta-analysis of AD:

AD QUERY A) [*Experiments Context IS Disease] AND [Experiment Contrast IS Gray Matter] AND [Experiments Observed Changes IS Controls>Patients] AND [Experiments Observed Changes IS Controls<Patients] AND [Subjects Diagnosis IS Alzheimer’s Disease]*;

AD QUERY B) [*Experiments Context IS Disease] AND [Experiment Contrast IS Gray Matter] AND [Experiments Observed Changes IS Controls>Patients] AND [Experiments Observed Changes IS Controls<Patients] AND [Subjects Diagnosis IS NOT Alzheimer’s Disease]*.

Therefore, two BrainMap taxonomy experts screened all the identified articles in order to ascertain that: a) a specific whole-brain VBM analysis was performed, b) a comparison between pathological sample and healthy control participants was included, c) GM decrease/increase changes in pathological sample were included, d) locations of GM changes were reported in a definite stereotactic brain space (i.e., Talairach or MNI).

On the basis of the aforementioned criteria, we included in our analyses: 114 articles, for a total of 147 experiments, 1754 GM changes and 4944 subjects (SCZ QUERY A); 693 articles, for a total of 1211 experiments, 9353 GM changes and 41746 subjects (SCZ QUERY B); 55 articles, for a total of 83 experiments, 961 GM changes and 1297 subjects (AD QUERY A); 760 articles, for a total of 1277 experiments, 10151 GM changes and 49194 subjects (AD QUERY B). Descriptive information of interest and meta-data of GM changes were extracted from each selected article (see Supplementary Table S1 for detailed information about the description and distribution of the VBM data set included in the meta-analysis). In order to facilitate subsequent analyses, coordinate-based results from MNI stereotactic space were converted into Talairach space by using Lancaster’s icbm2tal transformation (48, 49).

The selection of studies was performed according to the Preferred Reporting Items for Systematic Reviews and Meta-Analyses (PRISMA) Statement international guidelines (50, 51). The overview of the selection strategy is illustrated in Supplementary Figure S1 [PRISMA flow chart].

### Anatomical likelihood estimation and creation of the modeled activation maps

Although there are different methods of coordinate-based meta-analysis, the goal shared by all of them is to identify brain areas showing consistent activation across studies. These methods fall into two main approaches: kernel-based and model-based. The most important kernel-based methods are the multilevel kernel density analysis (MKDA) (52), the signed differential mapping (53), and the ALE (Eickhoff et al., 2012). The model-based methods use techniques derived from spatial statistics and develop stochastic model of analysis; among those the more developed are: the Bayesian hierarchical cluster process (54), the spatial binary regression model (55), the hierarchical Poisson/Gamma random field model (56), and the spatial Bayesian latent factor regression model (57). In all these models analyses are performed using Bayesian techniques in order to reconstruct spatial maps starting from lists of foci (spatial coordinates).

In our analysis we performed an ALE using the random effects algorithm of GingerAle (v. 2.3.6, http://brainmap.org) (Eickhoff et al., 2009; Eickhoff et al., 2012; Turkeltaub et al., 2012). The ALE is a quantitative voxel-based meta-analysis technique capable of providing information about the anatomical reliability of results through a statistical comparison on the basis of a sample of reference studies from the existing literature (58).

Each focus of every experiment is modeled by the ALE as a Gaussian probability distribution:

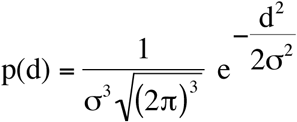

where d indicates the Euclidean distance between the voxels and the considered focus, and σ indicates the spatial uncertainty. Subsequently, we determined for every experiment a modeled alteration (MA) map as the union of the Gaussian probability distribution of each focus of the experiment. The union of these MA maps provided the final ALE map. The significance of the ALE activations are then tested against an empirical null distribution, thus yielding a P-value – see Eickhoff et al. (2012) for a detailed description of the algorithm. As the ALE creates smooth estimates of the local abundance of peak activations and is equivalent to a density estimation, for our purposes we normalized this density so as to obtain a probability function. To calculate the BF, however, we started from the unthresholded ALE maps (this choice was necessarily based on mathematical reasons; see the paragraph in which we discuss the transformation from ALE values to probability values).

### Reverse inference and Bayes’ factor

The framework of the reverse inference is the Bayes’ theorem. In general, reverse inference in neuroimaging provides information about the involvement of brain areas in cognitive processes. More precisely, making a reverse inference we can infer the likelihood of a specific cognitive process M from a certain pattern of brain activity A. As already said, the same reasoning still holds if we substitute “brain activity A” with “brain alteration A” and “cognitive process M” with “brain disorder M”. In this case, the reverse inference leads us to infer the posterior probability of a pathology P(M|A) from a certain pattern of brain alterations P(A|M) using the Bayes’ theorem:

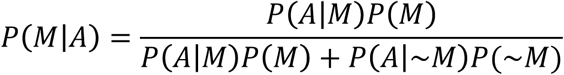

So, given the brain alteration A and the prior probability that the disease M occurs, it is possible to assess the posterior probability that M occurs on the basis of A.

The major issue that needs to be addressed in reverse inference is the determination of the prior probability. This choice depends on the information that we know before calculating the likelihood. In the literature, there is no general consensus as to how this choice should be made – for a review of the several proposals put forward to address this point see Carlin and Louis (59).

The Bayesian approach to testing hypotheses has been developed by Jeffreys (44) as part of his scientific program to study inference. Within this approach, given two competing hypotheses, the statistical models represent the probability that data are in accord with one or the other hypothesis and the Bayes’ theorem is used to determine the posterior probability of which of the two hypotheses is correct. In other words, if we assume that a set of data *D* can be accounted for by hypotheses *H*_1_ or *H*_2_, given a prior probability distribution *P*(*H*_1_) and *P*(*H*_2_) = 1 – *P*(*H*_1_), then according to the Bayes’ theorem the likelihood of each of the hypotheses to explain the data is:

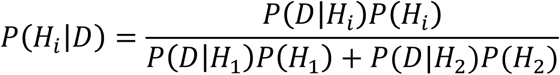

with *i* = 1,2. By calculating the ratio between the two hypotheses we have:

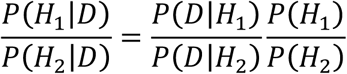

that can be expressed in words as follows:

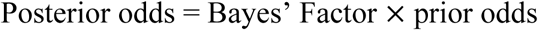

The BF is therefore the ratio between the posterior and prior odds, and represents a summary of evidence in favor of one of the two hypotheses. In our case the prior odds are set to 1 because we assume an equiprobable prior probability for each of the two hypotheses.

Our aim is to study the specificity and this can be obtained only and exclusively by considering the condition in which the alteration of the voxel is specific for the brain disorder under scrutiny.

So, by means of BF we compared the values of the MA maps related to each of the selected brain disorders (meta-data selected from BrainMap SCZ QUERY A and AD QUERY A) with the values of the MA maps related to all the other disorders (meta-data selected from BrainMap SCZ QUERY B and AD QUERY B). With this procedure, we determined a parameter of specificity capable of showing whether or not a certain brain area, if altered, was associated with a specific condition and, if so, how strong this association was. We were thus able to differentiate the ALE values associated with each of the diseases taken into account in this investigation.

For the analysis of VBM data the competing hypotheses were: i) the alteration of the voxel was specific for the brain disorder under scrutiny; ii) or the alteration was not (¬) specific. We needed therefore to calculate the ALE maps derived from all the experiments in which the detected alterations were associated with a specific brain disorder and the ALE maps derived from all the experiments in which the detected alterations were not associated with that specific brain disorder. To transform these ALE maps into a probability distribution map, we normalized the unthresholded maps so that the sum of all its values was equal to 1. The final result of this process was the BF, a number between [0, *∞*) representing how much the data could support the model *Disease* or ¬*Disease*. Following Kass and Raftery (47), the BF value can be interpreted as follows (Tab. 1):

**Table 1.**
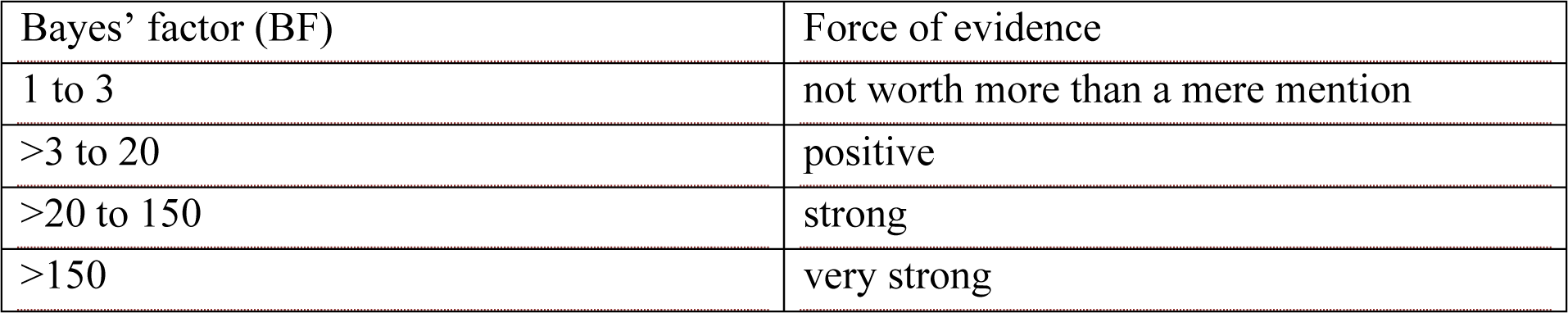
Bayes’ factor points associated with different forces of evidence.

So, if we have a BF of 4, which from the table indicates a positive evidence, it means that the hypothesis *H*_1_ is four times more probable than hypothesis *H*_2_. In our case, in which we used the posterior odd ratio, a BF of 4 means that the hypothesis that the detected gray matter alterations are associated with a specific brain disorder is four times more probable than the alternative hypothesis that these alterations are not specific for the pathology taken into account.

### Validation

To assess the efficacy of our algorithm we have performed an analysis of reverse inference on fMRI data about pain tasks obtained from the BrainMap database; this analysis has been already carried out by Yarkoni, Poldrack (28), and its results are also available on the Neurosynth platform (core tools) as well as in Yarkoni (39). We made the following two queries in the functional BrainMap database sector (April 2018):

A-PAIN) [*Experiments Context IS Normal Mapping] AND [Experiments Activation IS Activation Only] AND [Subjects Diagnosis IS Normals] AND [Experiments Imaging Modality IS fMRI] AND [Experiments Paradigm Class IS Pain Monitor/Discrimination]*;

B-NO PAIN) [*Experiments Context IS Normal Mapping] AND [Experiments Activation IS Activation Only] AND [Subjects Diagnosis IS Normals] AND [Experiments Imaging Modality IS fMRI] AND [Experiments Paradigm Class IS NOT Pain Monitor/Discrimination]*;

We retrieved 81 articles, for a total of 261 experiments, 2604 foci and 1157 subjects (QUERY A); and 3141 articles, for a total of 10209 experiments, 87409 foci and 58367 subjects (QUERY B) (see Supplementary Figure S2 [PRISMA flow chart] for the overview of the selection strategy, Table S2 for the sample characteristics and Table S3 for more information about the selected fMRI data set, respectively).

From the data retrieved with Sleuth 2.4 we calculated the ALE for the condition “pain” and for the condition “no-pain”. On the basis of the priors *p*(*H*_*i*_) = 0.5, *i* = 1,2, we determined the posterior probability and then the BF as the posterior odd ratio:

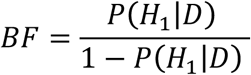

The results of our analysis were compared visually with those of Poldrack (23) as they are illustrated in Fig. 2 of his study (23) as well as with the results of an association map produced by Neurosynth (28). Indeed, the standard output of Neurosynth is not actually a BF map, so comparisons are necessarily to be made by analogy.

### Validation of the priors choice: Schwarz criterion and Bayes’ information criterion

The Bayes’ factor density *P*(*D*|*H*_*k*_) is obtained with an integration to the space of parameters, as follows:

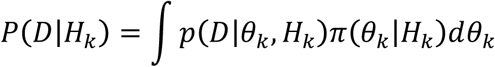

where *θ*_*k*_ are the parameters of the hypothesis *H*_*k*_ and *π*(*θ*_*k*_|*H*_*k*_) is the prior density, which depends on the assumed model of the data to be analyzed. Defining this prior is very difficult, unless the assumed model is deemed to be reliable. In fact, according to Yarkoni et al. (2011), with the empirical Bayes’ method it is not possible to derive from neuroimaging data a clear indication about the priors, unless these data are constituted by very large samples. However, it is possible to avoid the introduction in the model of the prior density with the help of the following formula:

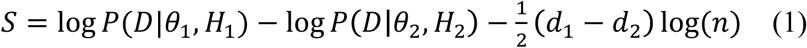

where *θ*_*k*_ are the estimations with regard to the different hypotheses *H*_*k*_, *d*_*k*_ is the dimension of *θ*_*k*_, and *n* is the numerosity of the sample. If *n ⟶ ∞*, it is possible to show the validity of the following relation, called Schwarz criterion (46):

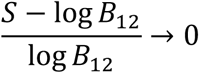

The Schwarz criterion, therefore, tends to the BF. If we consider the BIC, which is defined as:

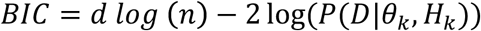

we can see that, from (1), minus twice the Schwarz criterion is the difference between the BIC of the two hypotheses – for an in-depth description of this theme see Kass and Raftery (47). Finally, if we multiply the BIC for minus 0.5 we obtain the S value, which can be compared with the BF. As pointed out in the introduction, we can observe that the BIC depends on the number of samples (log (*n*)); it is therefore a way of verifying whether or not the choice of 0.5 as prior is correct for the BF calculus. If we obtain convergent results, we would be relatively confident that the rationale for choosing equiprobable priors is, at least with large samples, sound.

### Stability against sample unbalances: Sample unbalance compensation

To reduce the potential bias of the uneven representativeness of brain disorders in the database, we created a single sample for each pathology by calculating an ALE map of all the studies related to a specific disease. BF maps constructed with this method were then correlated to other maps constructed without compensation.

### The file drawer problem or Robustness against noise: Fail-safe

The “fail-safe” technique is frequently used in classical meta-analyses of both medical and psychological studies. It was first introduced by Rosenthal (60) and further developed by Acar, Seurinck (61) for assessing the robustness of results against potential publication bias in coordinate-based fMRI meta-analyses. This method presumes that there are unpublished studies with significant as well as not significant results, and consequently estimates the number of these studies so as to modify the results of the meta-analysis. To do so, the method requires to introduce into the sample an amount of noise in order to assess how robust is the statistical result. Here the “fail-safe” technique has been used to address the possibility that in BrainMap an amount of significant and/or not significant experiments has not been stored, or that the different brain disorders are not uniformly represented in the database; this could be due to the fact that experiments about some disorders are more frequently stored than others and/or to the fact that certain pathological conditions are more investigated than others.

We used the algorithm proposed by Acar, Seurinck (61), which is available on Github (https://github.com/NeuroStat/GenerateNull). The procedure was the following: a vector reporting the data (i.e., number of foci and subjects for each experiment) is generated from the same list of foci used for the ALE, then the number of random experiments (i.e., noise value) is arbitrary set. The number of foci and the number of subjects of these random experiments are generated on the basis of the previous vector. For example, if the original meta-analysis consisted of foci between 10 and 20 and subjects between 10 and 20, the simulated experiments will also have a number of foci and subjects comprised between 10 and 20. The localization of peaks is uniformly and randomly sampled by a gray matter mask.

In order to assess the robustness of the data extracted from BrainMap we generated for each sample (i.e., AD, SCZ, and pain) a series of simulated studies between 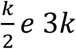, where *k* is the number of the original studies. These data were fed into the ALE and from these results we obtained the BF. In the simulation we used in GingerAle 1000 permutation for every noise value to test for the significance of the results. Subsequently, to analyze the results we used the following metric: a BF map (with a threshold level of 3 BF points) was obtained from the original studies and used as a mask for the analysis of the BF maps (with a threshold level of 3 BF points) obtained from the simulated studies. Following the method of comparison proposed by Acar, Seurinck (61), we calculated the correlation between these maps and saw its trend in relation to the number of simulated studies. Finally, for each sample we created a map showing the degree of alteration of each brain area after an increasing injection of noise.

### The Bayes’ factor and temporal evolution of brain diseases

We run a simulation to understand the capacity of BF to highlight the earliest areas to be altered. As already showed (18, 24, 62-66), neuropathological alterations are supposed to be distributed across the brain following structural and functional connectivity pathways. In order to simulate the alteration spread related to a certain pathology we used the anatomical connectivity matrix derived from Hagmann, Cammoun (67). Specifically, we used the average fiber tract density between two brain areas, obtained from healthy individuals with a parcellation of gray matter in 998 areas (nodes). First, we simulated a target pathology, that is, AD; AD is the ideal candidate because its areas of inception are well known. To do so, we selected 3 nodes for each cerebral hemisphere on the basis of the anatomopathological knowledge about AD (68). These 6 nodes were selected because of their proximity to the transentorhinal cortex, which is known to be one of the earliest sites of neurofibrillary deposition in AD (69), and were used as starting points for a simulated pathological spread. The model of simulation was based on the diffusion equation already applied in Cauda, Nani (24), which is the following:

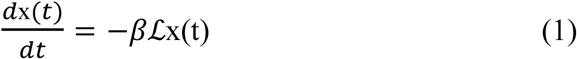

where the matrix *ℒ* is the following Laplacian graph:

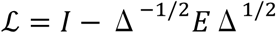

in which Δ is the diagonal matrix with *δ*_*i*_ = Σ_*j*_*e*_*ij*_as the *i*th diagonal element.

The solution of this equation is:

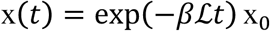

This formula describes the evolution of an initial configuration x_0_. The initial condition determines where the pathology begins to spread, that is, which nodes are the earliest to be altered. Thus, by considering different initial conditions, it is possible to simulate different temporal evolutions of brain diseases. The entire temporal span of the alteration spread was subdivided in 1000 time points (arbitrary units), from the initial condition (in which few nodes are altered) to the state of equilibrium (in which all the nodes are altered). After obtaining the diffusion data, we randomly selected for each simulation 100 time points so as to have a simulated picture of the uneven distribution of pathological alterations. In each of these time points we analyzed for every node the degree of its alteration; the nodes that showed a degree of alteration over a predetermined threshold were considered as being actually altered. Subsequently, every selected time point with its surviving altered nodes was treated as an experiment, thus generating different MA maps obtained from simulated patients (from 6 to 20); all these MA maps were eventually united in an ALE map.

To simulate other possible pathologies (up to 30), we selected for each disease 6 bilateral nodes (3 for each side) that were used to study the temporal evolution of the alteration spread. Overall, we generated 1000 time points for every simulated disease. As we did for the target pathology, we randomly selected 100 time points, so as to analyze the uneven distribution of alterations in different temporal series. Simulating different number of patients (from 6 to 20), each time point generated an MA map, which was united with the other MA maps of the other time points in an ALE map. Finally, the MA maps obtained from the target pathology (i.e., AD) and those obtained from the other simulated diseases were used for the BF calculus.

## Results

### Bayes’ factor

#### Comparison with previous results

To test our algorithm we replicated the analysis performed in Yarkoni, Poldrack (28). Fig. 1 (middle and bottom rows) shows the results of the reverse inference on pain tasks data queried from BrainMap as well as the association map provided for ‘pain’ by Neurosynth. Our results are very similar, albeit more conservative, to those of Neurosynth as well as to those shown by Yarkoni (28, 39). However, differences are to be expected, given the variability of input data and the methods used for constructing the maps (ALE vs MKDA; see (52). Moreover, Neurosynth does not provide a BF map but an association map expressed in z scores – for a detailed discussion see (28, 39).

**Figure 1.**
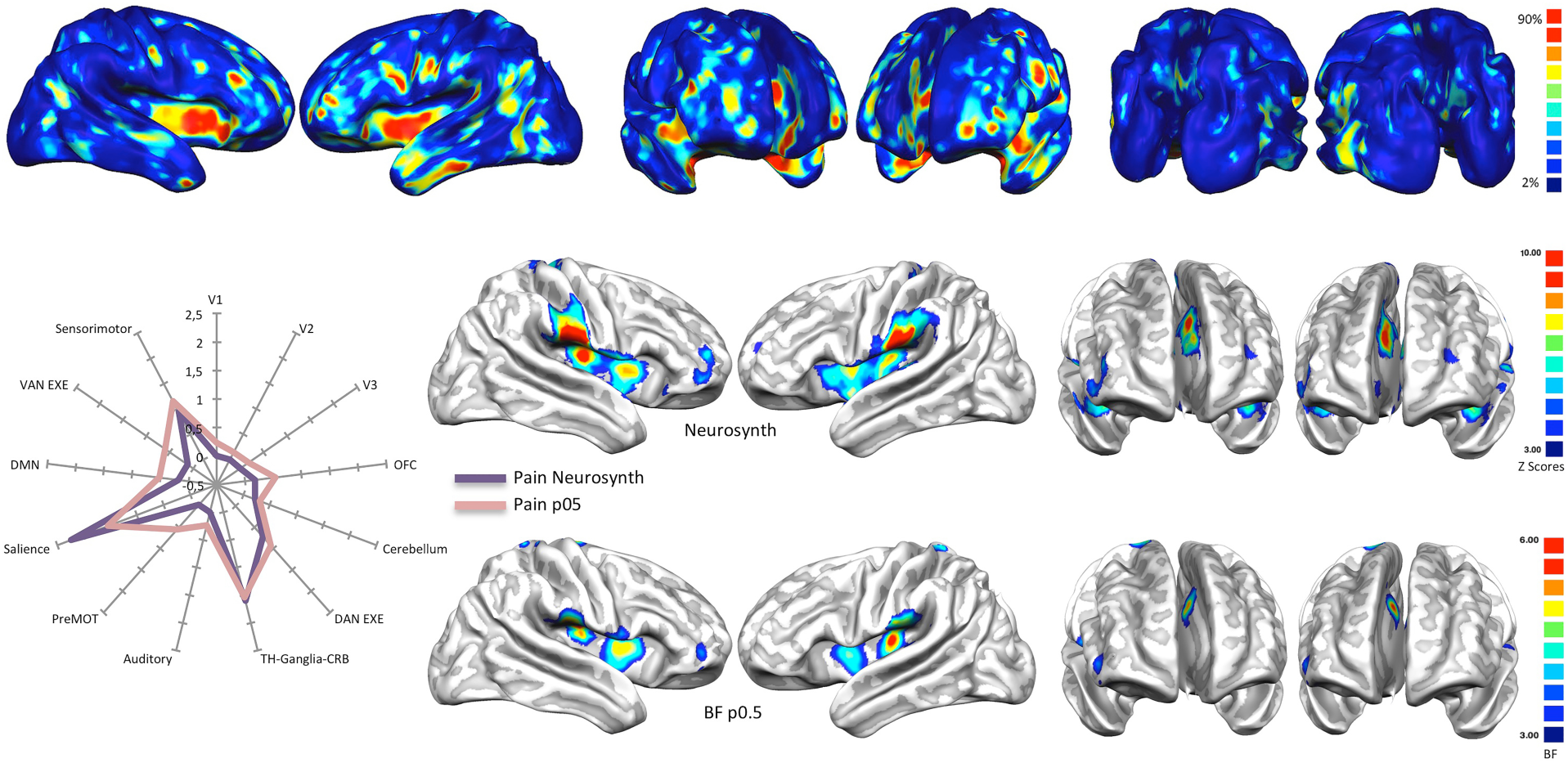
*Top:* Base rates reports of how many pathologies of the VBM BrainMap database cause alterations in every area of the brain. Areas highlighted in yellow/green are those in which more than the 90% of pathologies cause at least an alteration. *Middle and bottom*: Association test (expressed in z points) over the term ‘Pain’ performed with Neurosynth and compared with a Bayes’ factor map calculated with equiprobable priors over BrainMap data. *Left panel:* Radar map illustrating the comparison between the network-based decomposition of previous results expressed in z mean points (Neurosynth) and Bayes’ factor values.

#### Alzheimer’s disease and schizophrenia

The maps of specificity reveal that the two most represented pathologies in BrainMap (i.e., AD and SCZ) are characterized by certain areas with positive BF values (i.e., increased specificity). Most of the areas show positive (between 4 and 20) but not very strong BF values (>20) (see Fig. 2, top right and bottom right, and Supplementary Results in the Supplementary Material). Only few brain areas show strong BF values.

**Figure 2.**
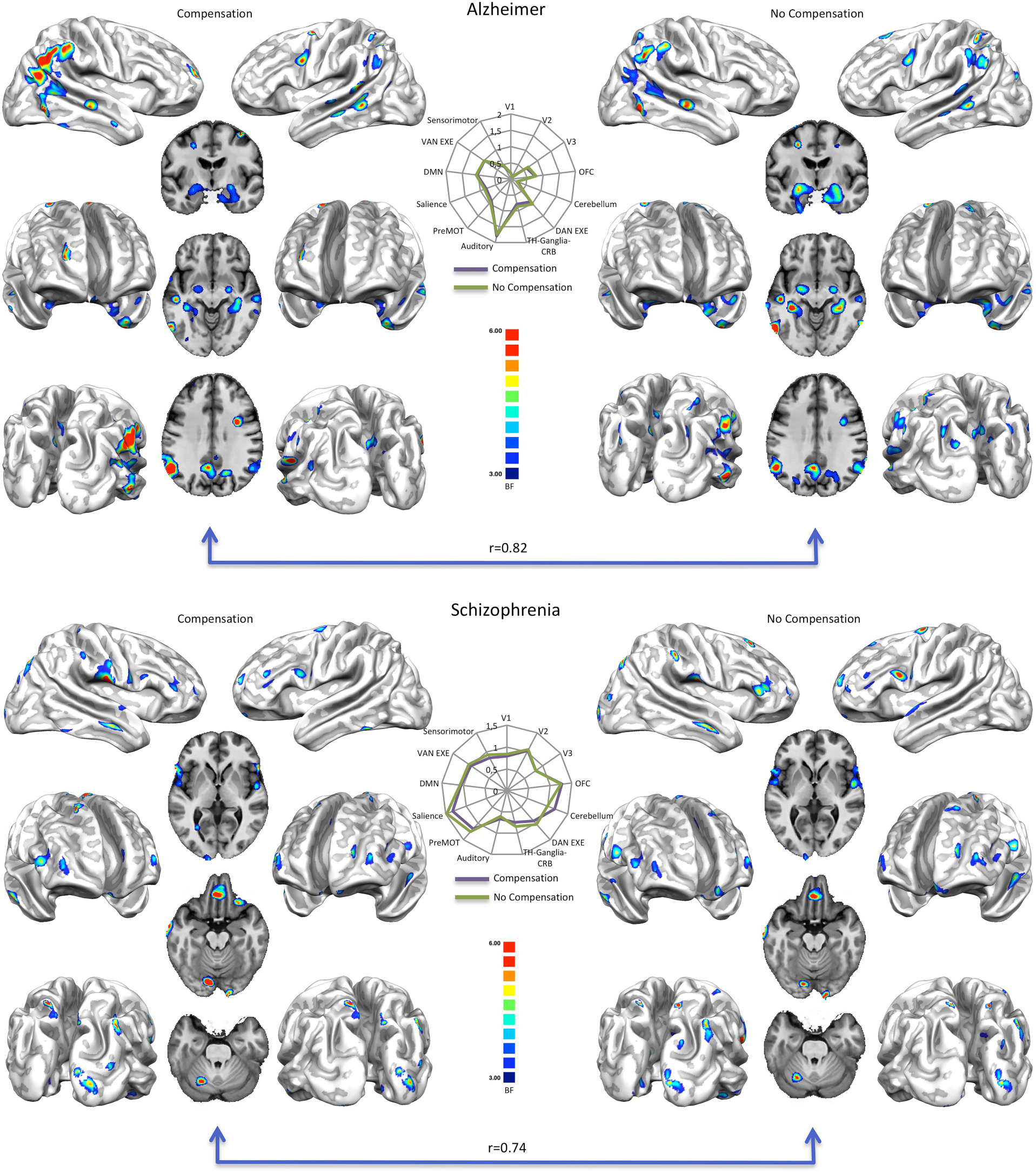
*Top right:* Bayes’ factor map of Alzheimer’s disease calculated with equiprobable priors over BrainMap data. *Top left:* Bayes’ factor map of Alzheimer’s disease calculated with equiprobable priors over BrainMap data, compensated for the different representativeness of pathologies in the database. *Bottom right:* Bayes’ factor map of schizophrenia calculated with equiprobable priors over BrainMap data. *Bottom left:* Bayes’ factor map of schizophrenia calculated with equiprobable priors over BrainMap data, compensated for the different representativeness of pathologies in the database. *Middle:* Radar maps illustrating the comparison between the network-based decomposition of previous results expressed in mean Bayes’ factor values.

Overall, most of the brain areas show BF values that are inferior to 4, save for a few regions exhibiting positive, albeit not strong, values. For SCZ, the postcentral gyrus, orbital, inferior and superior frontal and superior temporal areas show positive middle values (between 10 and 20) (Tab. S1). For AD, the parahippocampus has strong BF values (superior to 20), whereas middle values are exhibited by the inferior parietal lobule and the caudate tail (see Tab. 1 for a numeric visualization). The highest BF values in the parahippocampus and the transentorhinal regions (70) demonstrate that high BF values are telltale signs of the cerebral areas that are the earliest to be altered.

Strictly speaking, we cannot consider the areas with the highest BF values as utterly specific for a certain pathology, as we have to take into consideration the problems raised by the representativeness of the database (see the Limitations section for a discussion). Still, we can compare results obtained from different pathological processes with each other, as well as compare different patterns of GM alteration associated with the same pathology, so as to reach a pathological imprint that is more specific to that pathology. This reverse inference may allow us to better distinguish, among the areas that are altered by a certain pathology, the ones having higher or lower specificity.

#### Validation

In addition to the comparison between our results related to pain data and those produced by Neurosynth, we have employed the following validation techniques.

#### Sample unbalance compensation

To minimize the potential bias of the inhomogeneous representativeness of brain disorders in the database, we generated a single sample for every disease by determining an ALE map of all the experiments about a specific pathology. Subsequently, the BF maps obtained with this compensatory procedure (Fig. 2, top left and bottom left) were correlated to the original BF maps obtained without compensation (Fig. 2 top right and bottom right). Correlation values are very high: 0.82 for AD and 0.74 for SCZ, respectively (see Fig. 2, middle panel). The fact that maps with and without compensation are substantially similar leads us to think that the uneven representativeness of brain disorders in the database may cause a limited bias in the attribution of BF values to the altered brain areas.

#### Bayes’ information criterion

To test how much the choice of equiprobable priors can influence the BF values, we compared the BIC with the BF map related to pain and generated with equiprobable priors (Fig. 3). Results show that both the techniques produce maps that are extremely similar to each other (r = 0.88). This high correlation leads us to think that the choice of equiprobable priors does not bias the BF calculus.

**Figure 3.**
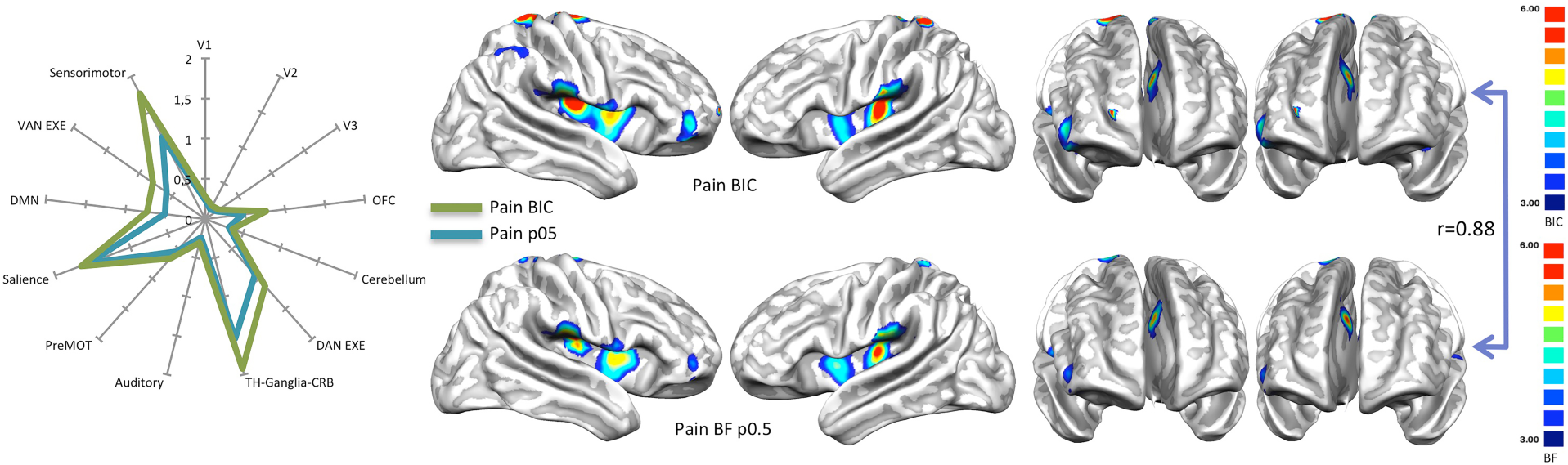
*Right panel:* Comparison between the results of the Bayes’ information criterion (BIC), expressed in S value (see Methods section), performed over the term ‘Pain’ and the Bayes’ factor (BF) map calculated with equiprobable priors over BrainMap data. *Left panel:* Radar map illustrating the comparison between the network-based decomposition of previous results expressed in mean BIC and BF values.

#### Fail-safe

To test the robustness of our results and evaluate the impact of the potential bias due to the heterogeneity of the database, we performed an analysis using the fail-safe technique. This procedure introduces progressively an amount of noise (i.e., studies with random results) with a percentage ranging from 25% to 300%. This analysis has been carried out with regard to AD, SCZ, and the pain condition. Results of the fail-safe analysis are illustrated by Figures 4 and 5.

Compared to the results related to AD and SCZ, the ones related to the pain condition are more vulnerable to the noise injections. In fact, for the pain condition most of brain areas with significant BF values do not survive after an amount of noise over 100% (Fig. 4, left panel). In this case, the most surviving areas are the sensorimotor and, to a lesser extent, the posterior insular. In contrast, AD and SCZ have higher correlations even with great quantities of noise injection. In particular, with regard to SCZ, the correlation is still at r = 0.67 after an amount of noise of 300%, while with regard to AD, the correlation is at r = 0.55 after an amount of noise over 150% (Fig. 4, middle and right panels). In the case of AD, the most surviving areas are those associated with the posterior component of the default mode network (DMN). In the case of SCZ, most of the areas with significant BF values survive after huge noise injections, save for the most anterior prefrontal areas. Figure 5 shows how the correlation between BF maps, derived from AD, SCZ, and the pain condition and obtained with and without noise, drops at r = 0.3 after an amount of noise over 150%.

**Figure 4.**
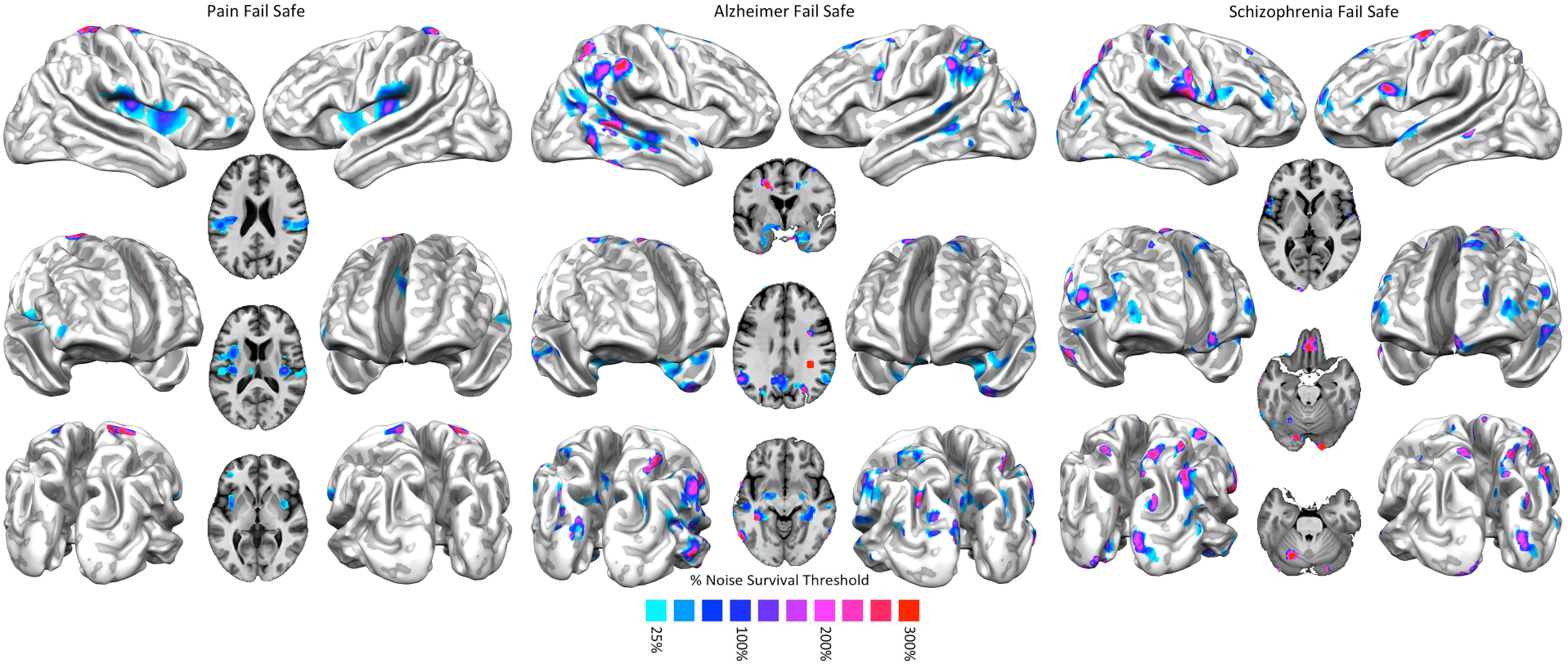
*Left panel:* Fail-safe results of the Bayes’ factor calculated over pain data. Areas colored from blue to red show increasing resistances to progressively greater injections of noise in the data set. *Middle panel:* Fail-safe results of the Bayes’ factor calculated over Alzheimer’s disease data. Areas colored from blue to red show increasing resistances to progressively greater injections of noise in the data set. *Right panel:* Fail-safe results of the Bayes’ factor calculated over schizophrenia data. Areas colored from blue to red show increasing resistances to progressively greater injections of noise in the data set.

**Figure 5.**
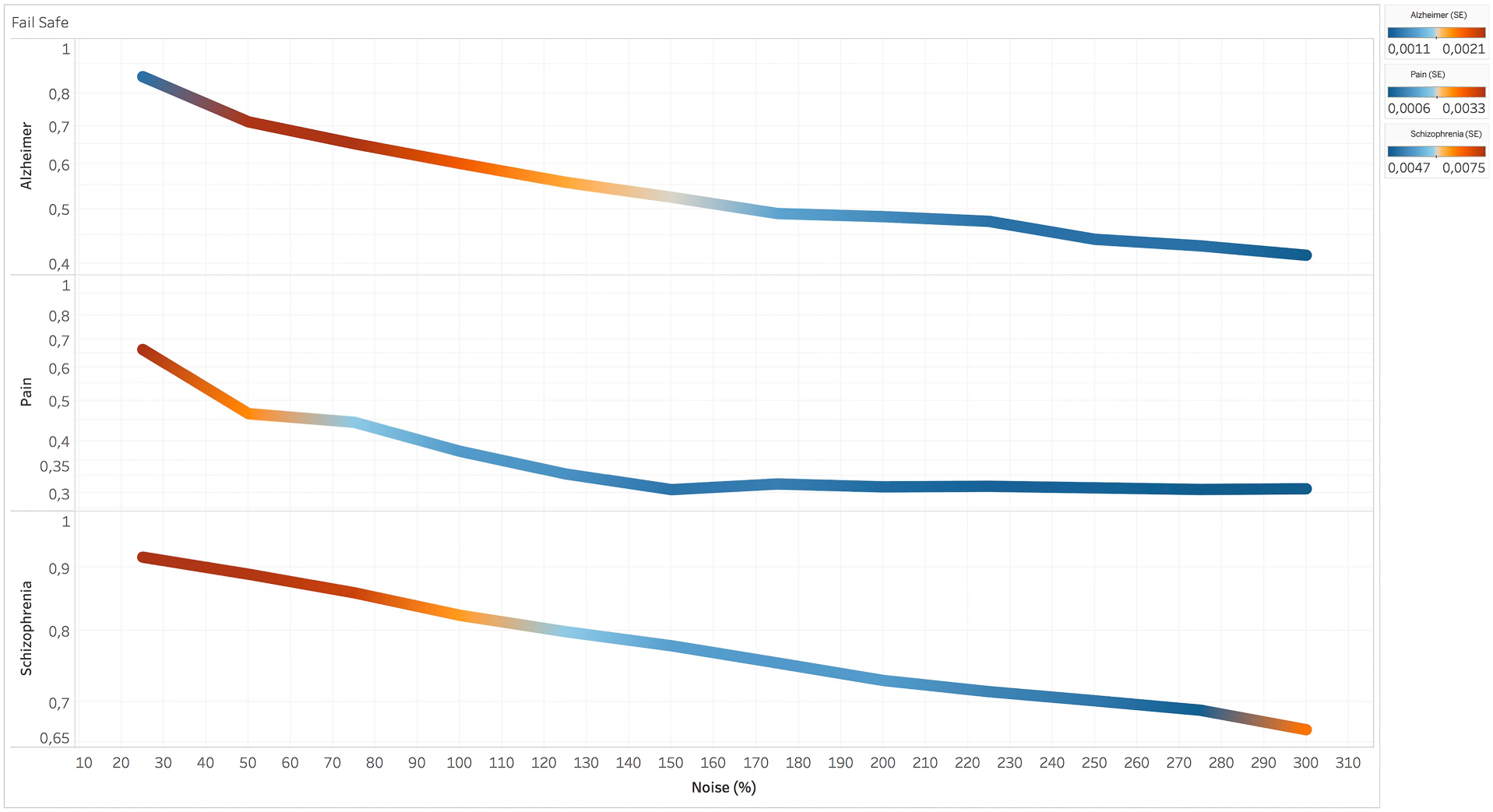
Fail-safe results of the Bayes’ factor (BF) of three data sets (i.e., pain, Alzheimer’s disease, and schizophrenia) obtained from the correlational values between the BF map calculated without injections of noise and BF maps calculated with progressively increasing injections of noise. Colors from blue to red indicate increasing values of standard deviation between the r values calculated in the different runs of the fail-safe procedure.

### The Bayes’ factor and temporal evolution of brain diseases

The simulation performed to demonstrate that the BF calculus is able to highlight the cerebral areas that are the earliest to be altered by a specific pathology can provide valuable insights in order to differentiate between areas precociously affected (more informative and specific) and areas that are altered later (less informative and specific) and, therefore, characterized by a greater overlap of alterations caused by different pathologies. Figure 6 shows the temporal evolution of the simulated target pathology (i.e., AD), the foci from which the pathology begins to spread (top left panel), and the temporal evolution of all the other simulated pathologies used for statistical comparison (middle panel). The figure also illustrates the BF map calculated on the synthetic data (bottom panel), as well as the comparison between the areas showing a BF>10 and the starting points (nodes) of the simulated target pathology (right panel). Clearly, our simulation shows that the BF calculus can capture, albeit with some approximation, the earliest points of the simulated spread. However, even if this Bayesian approach is not designed to precisely described the starting point of a specific disease, the conceptual implication suggested by high BF values deserves attention: in fact, the more specific is a brain area for a disease, the greater the likelihood for this area to be precociously involved in the progression of the disease. So, although the BF calculus cannot exactly pinpoint the very first starting site of a neuropathological process, it can identify the areas that are the earliest to be affected and differentiate them from those that appear to be affected later. Finally, the analysis of real data about AD has corroborated this approach, as it has revealed that the parahippocampus is the area with the strongest BF values (superior to 20; see Tab. 1 for a numeric visualization), a result that is in line with well-established scientific evidence (70).

**Figure 6.**
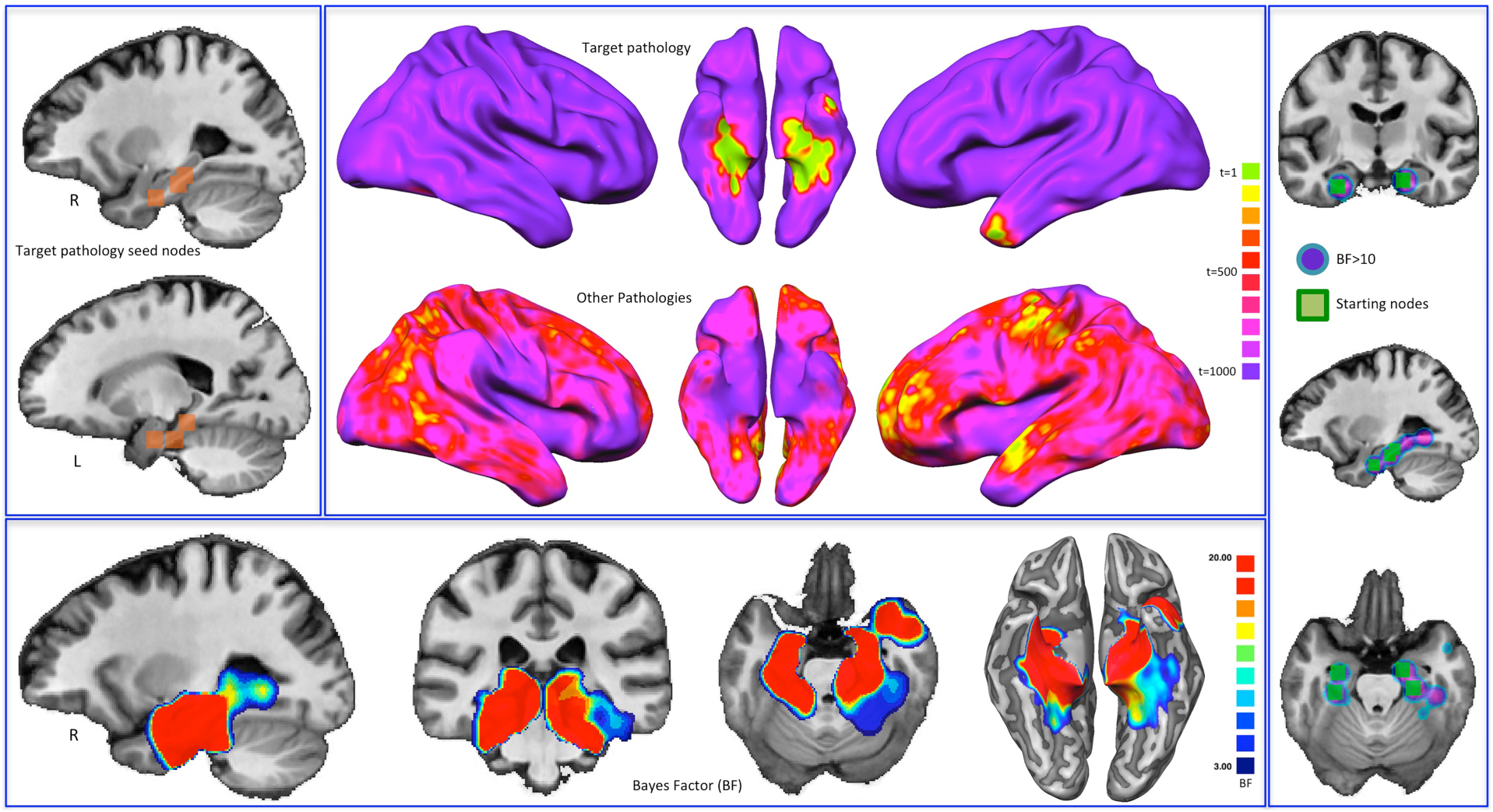
Bayes’ factor (BF) and the temporal evolution of pathologies. *Top left:* Starting nodes of the target pathology (Alzheimer’s disease, AD). *Middle:* Temporal evolution (expressed in arbitrary time points) both of the target pathology and of all the other simulated pathologies. Colors from green to violet show the areas that are altered from early to late phases of the simulated pathological spread. *Bottom:* BF values calculated on synthetic data. *Right panel:* Comparison between the areas showing a BF>10 and the starting points of the simulated target pathology.

#### Discussion

In this study we have applied a Bayesian reverse inference method to map the pathological brain and identify its altered regions that are specific to the two most represented disorders in BrainMap (i.e., AD and SCZ). This specificity is expressed in terms of BF, positive values of which (>3, but rarely superior to 20) characterize the structural alteration profiles both of AD and of SCZ. In particular, the posterior components of the DMN, the amygdalae, the hippocampus and parahippocampus exhibit positive specificity in the BF map of AD. Although the BF values of these areas are positive, they are not strong and just the parahippocampus shows a BF that is slightly superior to 20 (see Figure S3 and Tables S4, S5, and S6 in the Supplementary Material).

These findings are in accordance with well-established research and have been also further supported recently by our group (71). Volumetric changes involving the hippocampus/parahippocampus, and especially the entorhinal cortex, have been repeatedly considered as relevant features in the development of AD (72-74). In fact the atrophy of the medial temporal lobe, generally involving the amygdala (75), is a significant biomarker that helps predict the evolution from mild cognitive impairment to AD (76). Furthermore, resting-state fMRI investigations provide evidence that AD is associated with a decreased functional connectivity within the DMN (77). If we consider the progressive tauopathy that characterizes AD, the typical involvement of both entorhinal and parahippocampal districts may indicate that these are key regions for the deposition of pathologic tau in this condition (68, 78-82). Moreover, the deterioration of the mesial temporal (hippocampal/parahippocampal/entorhinal) cortex, in association with the posterior portions of the DMN, can be successfully identified by structural neuroimaging techniques (both VBM and cortical thickness) with different outcomes for sensibility/specificity in comparison to normal isocortical atrophy (83).

With regard to SCZ, we observe that the insula, anterior cingulate cortex (ACC), ventromedial prefrontal cortex, dorsolateral prefrontal cortex and medial thalamus show an increased specificity compared to other altered areas. The BF values of these areas are positive but not strong (ranging between 3 and 20). In morphometric studies of SCZ, the involvement of the insular cortex is frequently reported (84). Also, decreases in gray or white matter volumes have been observed in both ACC and various sites of the prefrontal cortex (85-92). The thalamus, too, is supposed to play a role in SCZ, on the basis of its many connections with several brain structures, especially with the prefrontal cortex (93, 94). The complex GM alterations, reported by scientific literature in patients with SCZ (in particular in insular and prefrontal regions) (95, 96), principally impact on areas whose disruption is directly associated with episodes of psychosis and their long-term outcome (97, 98), as well as with cognitive ability and the development of cognitive disturbances (99). Furthermore, the specific structural pattern, characterized by altered cortical thickness/volume associated with a perturbed cortical gyrification (both hyper-and hypogyria), reflects the involvement of a strong genetic or very early developmental background (100, 101). For all these reasons, SCZ can be considered as a model of neurodevelopmental disorder with high heritability.

The analysis of the alteration specificity put forward in this study is complementary to the transdiagnostic alteration patterns observed in previous investigations (4, 14-18, 23, 28, 29). These two approaches provide an overarching picture of the pathological brain, which is of great interest for a better understanding of how neuropathological processes affect this organ. Interestingly, although many cerebral areas are altered by the majority of brain diseases, patterns of alterations that seem specific to certain conditions can emerge. The study of these typical profiles of alterations promises to give valuable insights for the improvement of our clinical tools and for better understanding the development of brain disorders.

#### Validation

The analyses carried out in this study to understand how a sample that is not uniformly balanced, along with the choice for setting the priors, could bias the BF results have led us to think that the entity of the potential biases is not as serious as to invalidate the BF calculus. In fact the strategies used to overcome those difficulties, both in the case of the compensation of the inhomogeneity of the sample and in the case of the choice of the priors, have produced results very similar to those obtained without these procedures. Still, further research is needed to finally solve these problems. Especially with regard to the choice of the priors, we need further investigations with the help of empirical as well as Bayesian empirical techniques for their calculus.

The fail-safe analysis has showed that the BF results obtained from the anatomical alteration data of AD and SCZ are rather solid, as they survive after relevant injections of noise and they are not, therefore, strongly affected by publication bias (61). In particular, the results about SCZ are the best surviving, followed by the results about AD and, at a certain distance, by the results about the pain condition. Therefore, BF values obtained from VBM data were more resistant to publication bias than those obtained from functional data.

#### The temporal evolution of brain diseases

As already said in the introduction, it should be noted that when a specific brain disease (especially the neurodegenerative ones) is in its terminal stages, many areas of the brain will be affected. Therefore, if neuropathologies were studied during their advanced developments, they would show a great overlap of alterations. Differently from the discipline of pathological anatomy, which carries out post-mortem studies of the human brain, neuroimaging techniques usually examine mixed pathological populations exhibiting different phases of brain degeneration. Moreover, many studies are conducted on patients after their first diagnosis. These samples of individuals, therefore, are not expected to have areas showing great overlap of alterations (8).

Our simulations provide evidence that the BF calculus is able to identify the areas that are more precociously altered and, therefore, to distinguish between them and those regions that are affected later. This result emphasizes the importance to use reverse inference techniques for studying the anatomical alterations caused by brain diseases, as these techniques can provide a window into the initial stages of neuropathological development, thus making possible the identification of patterns of alteration spread as specific biomarkers of the early phases of brain disease, also in view of patients’ selection for disease-modifying clinical trials (102-104) as well as for assessing the efficacy of therapies (105-107). Indeed, with regard to AD and SCZ, with our Bayesian reverse inference method it has been possible to identify not only the areas that appear to be early affected (68, 108) but also the sites of the main pathological alterations of these two conditions, which are characterized by tauopathy and complex structural alterations related to a genetic/neurodevelopmental background (80, 100). This result supports well the hypothesis that high values of BF might be proportional to the degree of alteration earliness exhibited by those areas.

#### Limitations

The analysis on meta-analytical data, which are characterized by a certain degree of deterioration as well as of spatial uncertainty, might have increased the overlap between regions affected by different pathologies and, as a consequence, reduced the capacity of revealing small areas with high specificity. This aspect notwithstanding, the necessity of using a huge repository, in which data of a large number of brain disorders are stored, compels to adopt a meta-analytical approach. Although we have devised strategies to avoid the problem of the inhomogeneity of the BrainMap samples, and though the inhomogeneous distribution of data concerns the priors regarding the parameters of the likelihood model rather than the priors regarding the hypotheses, the uneven representation of pathologies may nonetheless have biased some results of the reverse inference. However, so far, the use of BrainMap is the only way to create maps of GM alterations capable of giving an overarching picture of the pathological brain. Indeed, as we have seen, a common described problem regarding reverse inference in fMRI is due to the inhomogeneous nature of the sample and to the estimation of the priors – see for instance (38, 39). As matter of fact, the frequency of distribution of pathologies in a database cannot represent accurately the frequency of distribution of pathologies in the real world. This is so because certain conditions are more investigated than others, independently of their incidence in the population, so that they are sampled with varying frequencies in the literature. Moreover, BrainMap contains only a fraction of all the neuroimaging literature about brain disorders and the addition of new studies is not principally directed to reduce this inhomogeneity. So, the amount of results that researchers report is inevitably related to their expectations as well as to the fact that some research topics are more prevalent than others; and the BrainMap database inevitably reflects that bias. As a consequence, our estimates may be biased upwards. Nonetheless, the validation analysis through the BIC, the compensation of the base rates of different pathologies and the fail-safe technique have showed that the potential biases of the sample are unlikely to invalidate our results.

Since a choice about the prior *must* be made, on the basis of our validation analyses and in absence of better strategies, we propose to choose equal priors (i.e., 50%). Needless to say, even though our validation analyses are encouraging, equal priors have both strengths and weaknesses; hopefully future procedures will provide better solutions. In the meantime, however, we think that the choice of equal priors is a valuable strategy and should be adopted. This would encourage the use of reverse inference techniques for the study of alterations caused by brain diseases, a study that, in our opinion, is essential to better understand the pathological brain, especially in light of the fact that many brain areas are non-specific to pathologies but, rather, exhibit a great overlap of alterations. Indeed, in spite of the problems already discussed, a study based on the reverse inference obtained from brain alterations data is extremely interesting, as it allows the identification of the areas that are more specific and/or precociously altered in a certain pathology. This obviously does not allow to sustain that the alteration of a certain area is strictly specific just to one pathology, but certainly it allows to identify which areas, whose alteration is frequently associated with more or less brain disorders, are more or less specific and informative.

Finally, our simulations of alteration spreads related to different pathologies are based on the premise that alterations move diffusively along brain connectivity pathways. Although this underlying mechanism has been confirmed by recent research (18, 24, 62, 63, 65, 66, 71, 109, 110), it is not the only one that might play a role in the alteration spread. Moreover, the contributions of different mechanisms can vary with regard to the type of pathology affecting the brain, so that our simulations, even though they offer in our view the best approximation to real pathological spreads with the available data, do not pretend to grasp all the complexities of the actual phenomenon.

A further limitation is that the BF is calculated in a univariate manner: each voxel or area is considered in isolation without taking into account a possible influence of other areas or voxels. Currently we are trying to develop an approach capable of considering a joint probability in which other variable as voxels or areas are taken into consideration; this view, however, poses several methodological problems, which are mostly related to the expansion of the parameters space and, consequently, to the difficulty in performing the calculation.

## Conclusion

Although transdiagnostic research provides evidence that many sites of the brain are altered by several pathological processes, this study shows that a Bayesian reverse inference is capable of identifying the cerebral areas exhibiting a high alteration specificity to certain pathologies. This approach allows to distinguish between areas that are altered by most of brain diseases and areas that are altered by a limited number of pathologies and can be, therefore, more specific to a certain pathology. It is also capable of identifying the areas that are the earliest to be affected or altered, thus opening a new window into the *in vivo* study of the pathological brain. These findings open interesting prospects for better characterizing brain disorders, as well as a new way to perform VBM meta-analyses, thus hopefully contributing to the intriguing quest for deciphering the complex landscape of alteration patterns of the pathological brain.

