## Supplementary online Materials for "Addressing reverse inference in structural brain alterations"

#### Supplementary Material


i. *Literature search strategy p. 2*

ii. *Supplementary Figures p. 4*

iii. *Supplementary Tables p. 7*

iv. *Supplementary Results p. 13*

### Literature search strategy

***Selection of Studies***

Neuroimaging experiments of interest were collected from the BrainMap database (http://brainmap.org/) (Fox and Lancaster, 2002; Laird *et al.*, 2005; Vanasse *et al.*, 2018). BrainMap is an online open access database of published functional and voxel-based morphometry (VBM) experiments with coordinate-based results in MNI or Talairach (TAL) stereotaxic brain space (Fox *et al.*, 2005). Notably, we used the software application 'Sleuth 2.4' to search the eligible meta-data and view the search results in MNI or TAL space.

As a first step, we queried the VBM BrainMap sector (Vanasse *et al.*, 2018) in order to retrieve the voxel-based results of the two most represented brain disorders of the VBM data set (i.e., schizophrenia (SCZ) and Alzheimer's disease (AD)). Therefore, we employed a double systematic search. The search algorithms were constructed as follows:

For the meta-analysis of SCZ (VBM sector):

a) [*Experiments Context IS Disease] AND [Experiment Contrast IS Gray Matter] AND [Experiments Observed Changes IS Controls>Patients] AND [Experiments Observed Changes IS Controls<Patients] AND [Subjects Diagnosis IS Schizophrenia]*;

b) [*Experiments Context IS Disease] AND [Experiment Contrast IS Gray Matter] AND [Experiments Observed Changes IS Controls>Patients] AND [Experiments Observed Changes IS Controls<Patients] AND [Subjects Diagnosis IS NOT Schizophrenia].*

For the meta-analysis of AD (VBM sector):

AD QUERY A) [*Experiments Context IS Disease] AND [Experiment Contrast IS Gray Matter] AND [Experiments Observed Changes IS Controls>Patients] AND [Experiments Observed Changes IS Controls<Patients] AND [Subjects Diagnosis IS Alzheimer's Disease]*;

AD QUERY B) [*Experiments Context IS Disease] AND [Experiment Contrast IS Gray Matter] AND [Experiments Observed Changes IS Controls>Patients] AND [Experiments Observed Changes IS Controls<Patients] AND [Subjects Diagnosis IS NOT Alzheimer's Disease].*

Finally, we made the following two queries in the functional BrainMap database sector:

A PAIN) [*Experiments Context IS Normal Mapping] AND [Experiments Activation IS Activation Only] AND [Subjects Diagnosis IS Normals] AND [Experiments Imaging Modality IS fMRI] AND [Experiments Paradigm Class IS Pain Monitor/Discrimination]*;

B NO PAIN) [*Experiments Context IS Normal Mapping] AND [Experiments Activation IS Activation Only] AND [Subjects Diagnosis IS Normals] AND [Experiments Imaging Modality IS fMRI] AND [Experiments Paradigm Class IS NOT Pain Monitor/Discrimination].*

***Assessment of eligibility of studies***

Two BrainMap taxonomy experts evaluated all the VBM meta-data identified by our search algorithms, thus ensuring that they comply with specific eligibility criteria. Articles were included if:

a) a specific whole-brain VBM analysis was performed;
b) a comparison between pathological sample and healthy control participants was included,
c) gray matter (GM) decrease/increase changes in pathological sample were included;
d) locations of GM changes were reported in a definite stereotaxic brain space (i.e, Talairach or MNI).

In case of fMRI meta-data, articles were included if:

a) a specific whole-brain fMRI analysis was performed;
b) the experimental sample corresponded to healthy participants;
c) locations of GM activation were reported in a definite stereotaxic brain space (i.e, Talairach or MNI).

Transparent and complete reports of VBM and fMRI data selection are shown in Figure S1 and Figure S2 [PRISMA flow chart] (Liberati *et al.*, 2009; Moher *et al.*, 2009).

***Data extraction***

We used the software application 'Sleuth 2.4' to extract the VBM and fMRI meta-data from each selected article. In order to facilitate our analyses, we converted the coordinate-based results from MNI space into TAL space by using Lancaster's icbm2tal transformation (Lancaster *et al.*, 2007; Laird *et al.*, 2010).

More information about the description of the extracted VBM data are viewable in Table S1. For more information about the selected fMRI data, see Table S2 and S3.

#### *References*


Fox PT, Laird AR, Fox SP, Fox PM, Uecker AM, Crank M*, et al.* BrainMap taxonomy of experimental design: description and evaluation. Human brain mapping 2005; 25(1): 185-98.

Fox PT, Lancaster JL. Opinion: Mapping context and content: the BrainMap model. Nature reviews Neuroscience 2002; 3(4): 319-21.

Goedert M, Masuda-Suzukake M, Falcon B. Like prions: the propagation of aggregated tau and α-synuclein in neurodegeneration. Brain : a journal of neurology 2017; 140(2): 266-78.

Laird AR, Lancaster JL, Fox PT. BrainMap: the social evolution of a human brain mapping database. Neuroinformatics 2005; 3(1): 65-78.

Laird AR, Robinson JL, McMillan KM, Tordesillas-Gutierrez D, Moran ST, Gonzales SM*, et al.* Comparison of the disparity between Talairach and MNI coordinates in functional neuroimaging data: validation of the Lancaster transform. NeuroImage 2010; 51(2): 677-83.

Lancaster JL, Tordesillas-Gutierrez D, Martinez M, Salinas F, Evans A, Zilles K*, et al.* Bias between MNI and Talairach coordinates analyzed using the ICBM-152 brain template. Human brain mapping 2007; 28(11): 1194-205.

Liberati A, Altman DG, Tetzlaff J, Mulrow C, Gotzsche PC, Ioannidis JP*, et al.* The PRISMA statement for reporting systematic reviews and meta-analyses of studies that evaluate health care interventions: explanation and elaboration. Journal of clinical epidemiology 2009; 62(10): e1-34.

Moher D, Liberati A, Tetzlaff J, Altman DG. Preferred reporting items for systematic reviews and meta-analyses: the PRISMA statement. Journal of clinical epidemiology 2009; 62(10): 1006-12.

Vanasse TJ, Fox PM, Barron DS, Robertson M, Eickhoff SB, Lancaster JL*, et al.* BrainMap VBM: An environment for structural meta-analysis. Human brain mapping 2018; 39(8): 3308-25.

### Supplementary Figures

**Figure S1 [PRISMA flow chart]**. Overview of the selection strategy (voxel-based morphometry data set). SCZ = schizophrenia; AD = Alzheimer's disease.

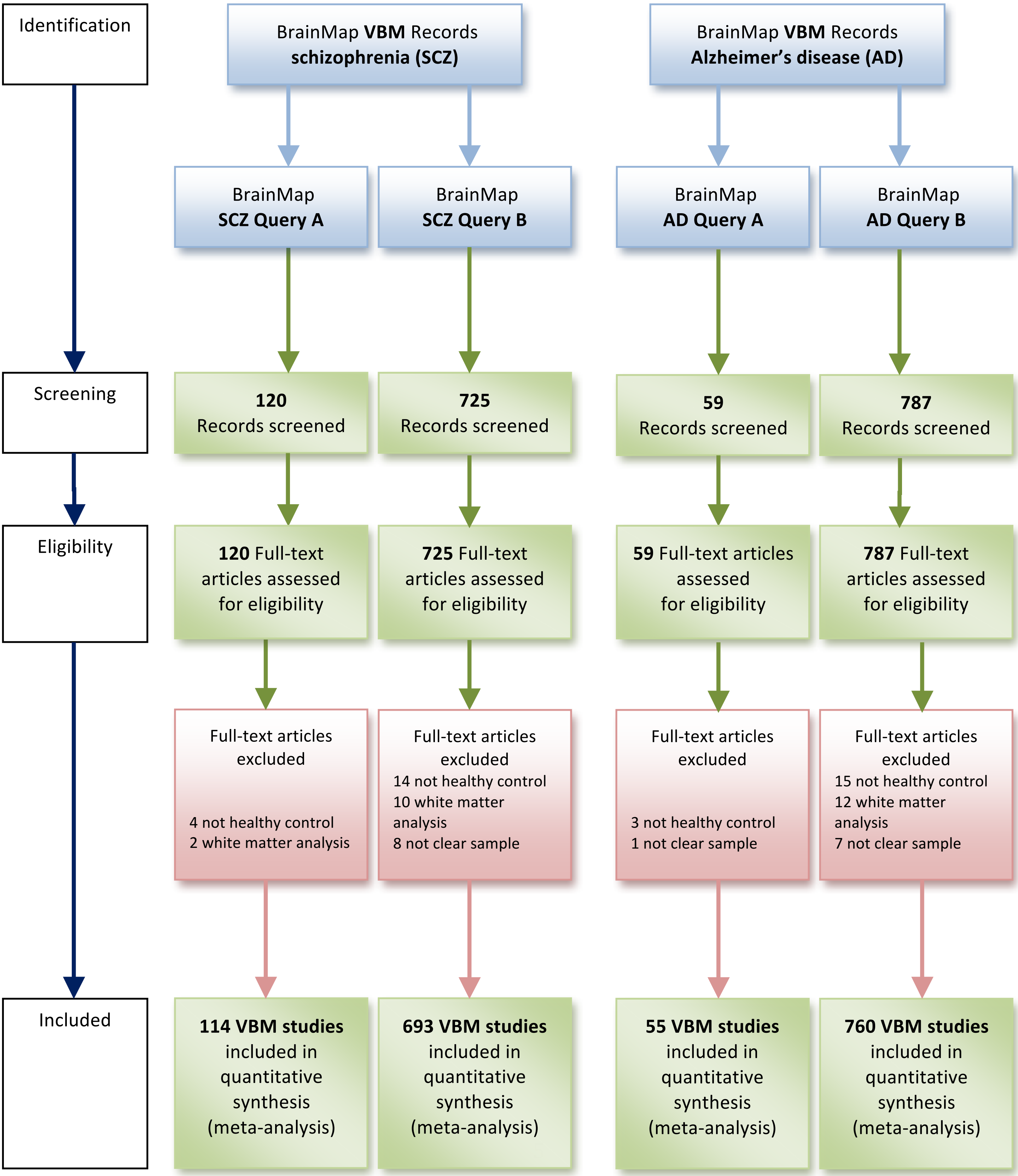

**Figure S2 [PRISMA flow chart]**. Overview of the selection strategy (fMRI data set).

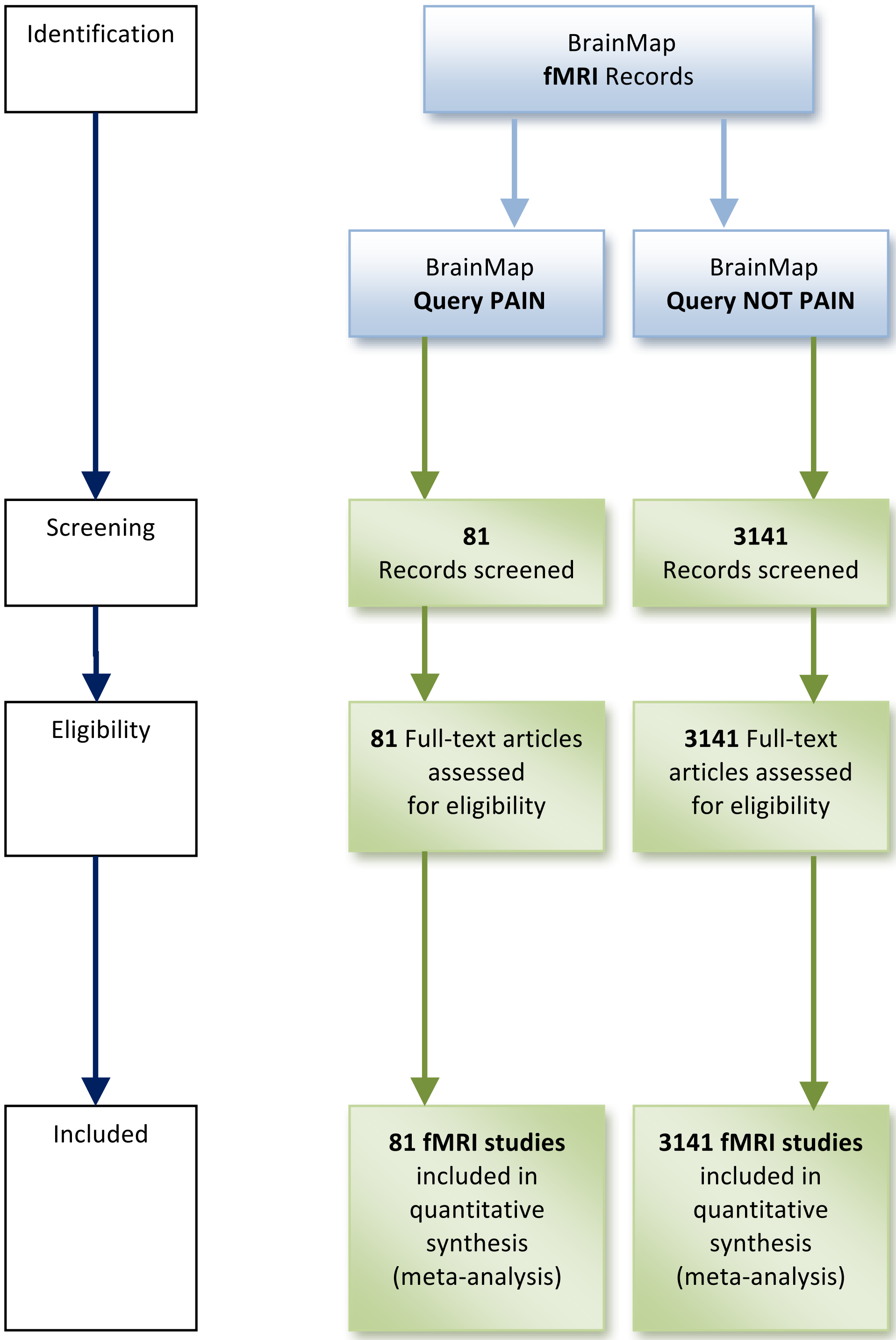

**Figure S3.** Graph reporting the Bayes' factor values for each cluster of Table S4 (schizophrenia), Table S5 (Alzheimer's disease) and Table S6 (pain).

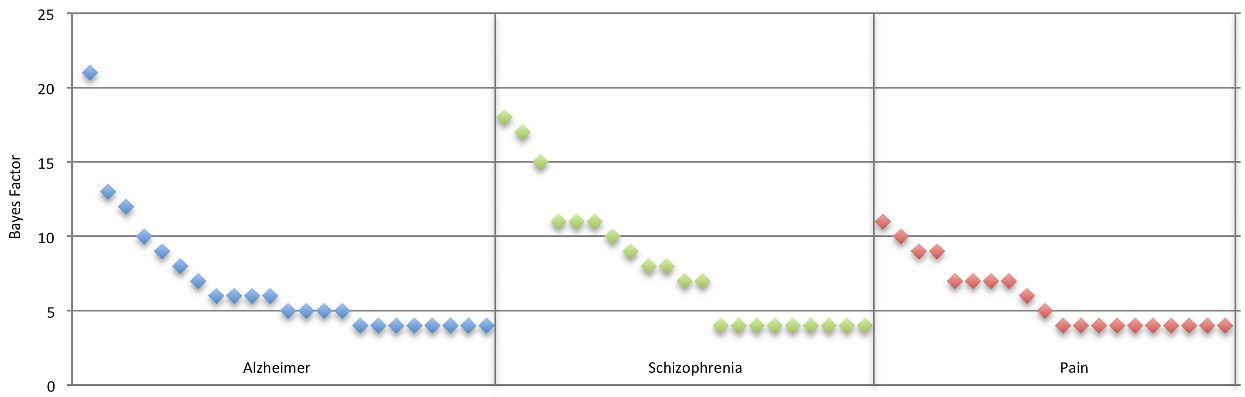

**Figure S4.** Infographic depicting the process through which the Bayes' factor can identify the early affected areas of a certain pathology. In this case the comparison between tau inclusion and the other two neurodegenerative aggregations provides evidence that the locus coeruleus is one of the first affected areas.
Image inspired by Michel Goedert, Masami Masuda-Suzukake and Benjamin Falcon, "Like prions: the propagation of aggregated tau and a-synuclein in neurodegeneration" (2017).

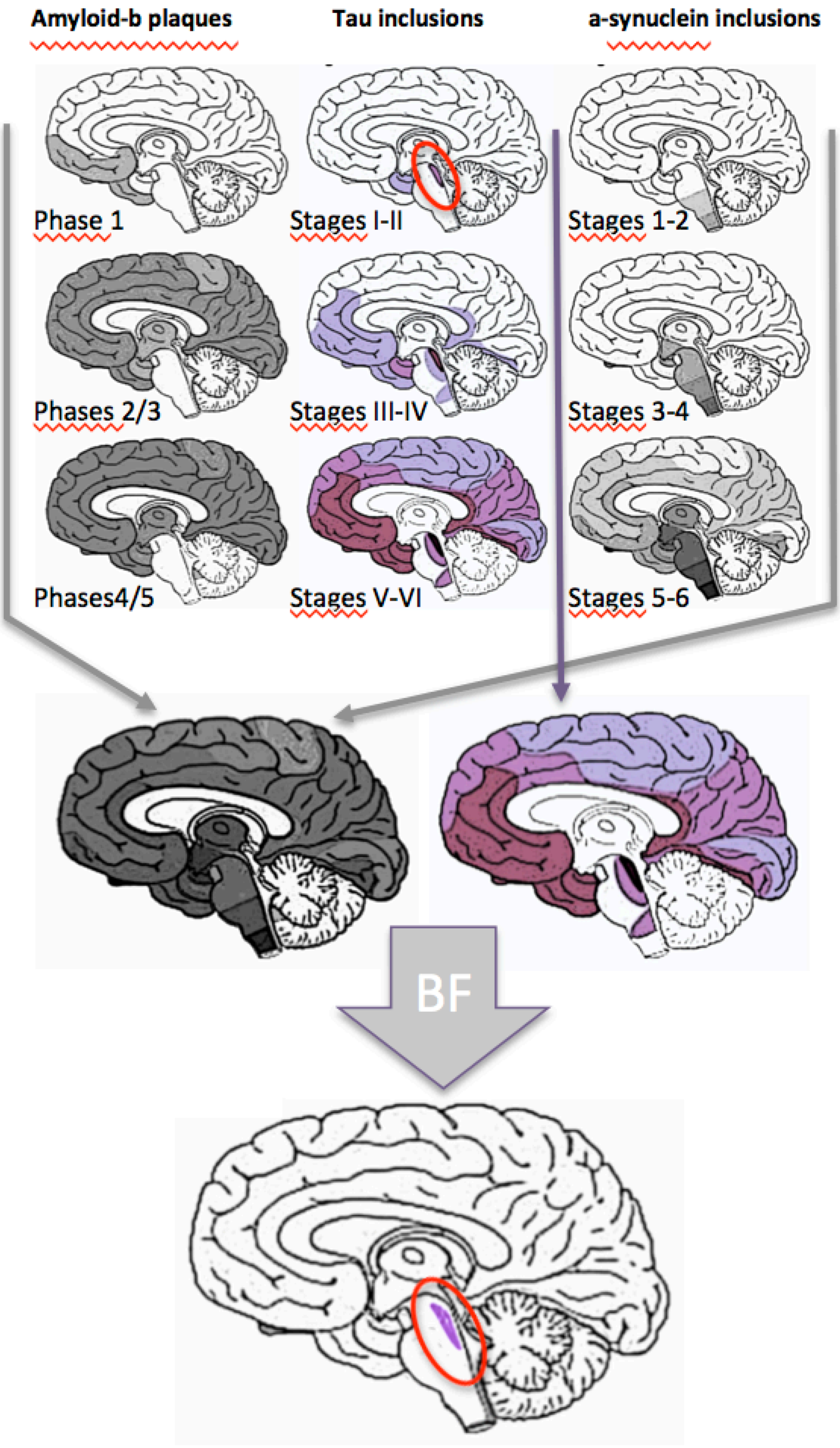

### Supplementary Tables

**Table S1**. Sample characteristics and gray matter (GM) variations with relative numbers of foci for each of the four queries made in the BrainMap voxel-based morphometry database.
Experiments (N) = number of experiments; Experiments (%) = percentage of the total of the selected experiments; Subj (N) = number of subjects.

| ***Query (Condition)*** | ***Articles*** | | ***Experiments*** | | ***Subj (N)*** | ***GM Changes*** | |
|---|---|---|---|---|---|---|---|
| | ***Decrease*** | ***Increase*** | ***(N)*** | ***(%)*** | | ***Decrease*** | ***Increase*** |
| SCZ QUERY A (IS SCZ) | 83 | 31 | 147 | 5.5 | 4944 | 1481 | 273 |
| SCZ QUERY B (IS NOT SCZ) | 513 | 180 | 1211 | 44.6 | 41746 | 7190 | 2163 |
| AD QUERY A (IS AD) | 44 | 11 | 83 | 3.0 | 1297 | 818 | 143 |
| AD QUERY B (IS NOT AD) | 559 | 201 | 1277 | 46.9 | 49194 | 7853 | 2298 |
| ***TOTAL*** | ***1199*** | ***423*** | ***2718*** | ***100*** | ***97181*** | ***17342*** | ***4877*** |

**Table S2**. Sample characteristics with relative numbers of foci for each of the two queries made in the BrainMap functional database.
Experiments (N) = number of experiments; Experiments (%) = percentage of the total of the selected experiments; Subj (N) = number of subjects.

| *Query (Condition)* | *Articles* | | *Experiments* | | *Subj (N)* | *Foci (N)* |
|---|---|---|---|---|---|---|
| | *(N)* | *(%)* | *(N)* | *(%)* | | |
| A (PAIN) | 81 | 2.5 | 261 | 2.5 | 1157 | 2604 |
| B (NO PAIN) | 3141 | 97.5 | 10209 | 97.5 | 58367 | 87409 |
| ***TOTAL*** | ***3222*** | ***100*** | ***10470*** | ***100*** | ***59524*** | ***90013*** |

**Table S3**. Types of paradigm classes with relative number of experiments, foci and subjects for each paradigm class of the fMRI BrainMap data set used in the meta-analysis (QUERY PAIN and QUERY NO PAIN).
Exp (N) = number of experiments; Subj (N) = number of subjects.

| ***Paradigm Class*** | ***Articles*** | ***Exp (N)*** | **Foci (x,y,z)** | ***Subj (N)*** |
|---|---|---|---|---|
| Acupuncture | 11 | 30 | 350 | 175 |
| Affective pictures | 50 | 175 | 1608 | 1182 |
| Affective words | 14 | 62 | 422 | 398 |
| Anti-saccades | 9 | 17 | 237 | 125 |
| Chewing/swallowing | 21 | 81 | 562 | 355 |
| Classical conditioning | 24 | 83 | 468 | 341 |
| Competition/cooperation | 5 | 29 | 188 | 116 |
| Counting/calculation | 56 | 170 | 1622 | 858 |
| Cued explicit recognition/recall | 78 | 250 | 2478 | 1374 |
| Deception | 18 | 49 | 521 | 310 |
| Delay discounting | 9 | 42 | 596 | 262 |
| Delayed match to sample | 75 | 224 | 2611 | 1070 |
| Divided auditory attention | 3 | 12 | 99 | 61 |
| Drawing | 2 | 15 | 78 | 20 |
| Driving | 5 | 15 | 218 | 60 |
| Emotion induction | 115 | 414 | 3021 | 2498 |
| Emotional body language perception | 4 | 15 | 163 | 55 |
| Encoding | 63 | 189 | 1654 | 1246 |
| Episodic recall | 21 | 76 | 669 | 311 |
| Estimation | 6 | 13 | 124 | 110 |
| Face monitoring/discrimination | 141 | 460 | 3609 | 2507 |
| Figurative language | 11 | 33 | 217 | 160 |

| | | | | |
|---|---|---|---|---|
| Film viewing | 60 | 211 | 2321 | 1046 |
| Finger tapping/button press | 287 | 1042 | 8389 | 6564 |
| Fixation | 16 | 36 | 153 | 137 |
| Flanker | 10 | 37 | 292 | 214 |
| Flashing checkerboard | 5 | 29 | 115 | 185 |
| Flexion/extension | 47 | 146 | 1356 | 636 |
| Fluency induction | 1 | 3 | 8 | 12 |
| Free list word record | 1 | 3 | 13 | 15 |
| Gambling | 34 | 151 | 1015 | 663 |
| Go/No go | 82 | 242 | 1798 | 2852 |
| Grasping | 10 | 30 | 229 | 111 |
| Hand-Eye coordination | 1 | 5 | 43 | 20 |
| Hunger/satiety | 8 | 22 | 77 | 114 |
| Hypercapnia/air hunger | 6 | 8 | 130 | 60 |
| Imagined movement | 28 | 85 | 905 | 476 |
| Imagined objects/scenes | 31 | 102 | 855 | 558 |
| Induced panic | 1 | 1 | 4 | 6 |
| Isometric force | 5 | 17 | 289 | 58 |
| Lexical decision | 11 | 26 | 193 | 165 |
| Magnitude comparison (distance) | 2 | 15 | 153 | 38 |
| Magnitude comparison (luminance) | 2 | 8 | 29 | 30 |
| Magnitude comparison (numerical) | 2 | 4 | 16 | 24 |
| Magnitude comparison (physical size) | 4 | 15 | 65 | 78 |
| Magnitude comparison (symbolic) | 6 | 16 | 123 | 99 |
| Meditation | 18 | 80 | 391 | 424 |
| Mental rotation | 26 | 94 | 765 | 416 |
| Micturition | 6 | 25 | 145 | 65 |
| Motor learning | 1 | 1 | 23 | 3 |
| Multi tasking | 5 | 14 | 191 | 82 |
| Music comprehension | 52 | 134 | 1930 | 978 |

| | | | | |
|---|---|---|---|---|
| Music production | 24 | 70 | 760 | 354 |
| N-back | 92 | 238 | 2175 | 2413 |
| Naming (covert) | 18 | 61 | 523 | 227 |
| Naming (overt) | 14 | 53 | 371 | 186 |
| Object manipulation/discrimination | 1 | 2 | 8 | 10 |
| Oddball discrimination | 21 | 48 | 605 | 376 |
| Olfactory monitoring/discrimination | 18 | 43 | 364 | 229 |
| Orthographic discrimination | 36 | 114 | 831 | 574 |
| Pain monitor/discrimination | 81 | 261 | 2604 | 1157 |
| Paired associate recall | 25 | 77 | 761 | 445 |
| Passive listening | 59 | 181 | 1402 | 890 |
| Passive viewing | 106 | 295 | 2966 | 1760 |
| Phonological discrimination | 51 | 173 | 1228 | 706 |
| Pitch monitor/discrimination | 30 | 118 | 1010 | 455 |
| Pointing | 6 | 15 | 145 | 63 |
| Pursuit rotor/manual tracking | 5 | 19 | 179 | 82 |
| Reading (covert) | 49 | 165 | 1510 | 713 |
| Reading (overt) | 23 | 79 | 1376 | 367 |
| Reasoning/problem solving | 53 | 173 | 1299 | 1161 |
| Recitation/repetition (covert) | 14 | 25 | 201 | 179 |
| Recitation/repetition (overt) | 22 | 55 | 799 | 266 |
| Rest | 7 | 17 | 140 | 128 |
| Reward | 168 | 700 | 5273 | 3302 |
| Saccades | 40 | 97 | 1014 | 469 |
| Self-reflection | 4 | 24 | 120 | 87 |
| Semantic monitor/discrimination | 153 | 513 | 3990 | 2579 |
| Sequence recall/learning | 20 | 60 | 625 | 306 |
| Sexual arousal/gratification | 18 | 64 | 991 | 437 |
| Sleep | 1 | 2 | 5 | 40 |
| Stroop - color | 41 | 88 | 889 | 869 |

| | | | | |
|---|---|---|---|---|
| Stroop - counting | 7 | 14 | 83 | 105 |
| Stroop - emotional | 11 | 21 | 124 | 301 |
| Stroop - other | 5 | 14 | 134 | 103 |
| Stroop - spatial | 3 | 10 | 71 | 36 |
| Syntactic discrimination | 11 | 18 | 126 | 191 |
| Tactile monitor/discrimination | 35 | 104 | 761 | 444 |
| Task switching | 37 | 117 | 1021 | 644 |
| Taste | 28 | 108 | 612 | 605 |
| Theory of mind | 54 | 228 | 1079 | 1787 |
| Thirst induction | 1 | 1 | 4 | 11 |
| Tone monitor/discrimination | 37 | 93 | 1084 | 555 |
| Tower of London | 4 | 6 | 77 | 60 |
| Transcranial magnetic stimulation | 3 | 8 | 96 | 32 |
| Trauma recall | 3 | 3 | 46 | 39 |
| Vestibular stimulation | 2 | 2 | 28 | 23 |
| Vibrotactile monitor/discrimination | 7 | 71 | 93 | 105 |
| Video games | 6 | 20 | 107 | 111 |
| Visual motion | 3 | 5 | 49 | 44 |
| Visual object identification | 42 | 146 | 1145 | 667 |
| Visual pursuit/tracker | 27 | 86 | 818 | 331 |
| Visuospatial attention | 91 | 270 | 2580 | 1337 |
| Wisconsin card sorting test | 13 | 37 | 286 | 237 |
| Word generation (covert) | 49 | 126 | 1169 | 650 |
| Word generation (overt) | 19 | 50 | 470 | 323 |
| Word imageability | 4 | 7 | 55 | 63 |
| Word stem completion (covert) | 4 | 6 | 57 | 64 |
| Word stem completion (overt) | 4 | 7 | 76 | 65 |
| Writing | 3 | 6 | 67 | 38 |
| ***TOTAL*** | ***3222*** | ***10470*** | ***90013*** | ***59524*** |

### Supplementary Results

**Table S4.** Clusters of specificity in schizophrenia (SCZ).
BF Value = Bayes' Factor Value

| *BF Value* | *Cluster* | *x* | *y* | *z* | *Side* | *Label* | *BA* |
|---|---|---|---|---|---|---|---|
| 18 | 2 | 56 | -18 | 18 | R | Postcentral Gyrus | 40 |
| 17 | 2 | 55 | -18 | 20 | R | Postcentral Gyrus | 43 |
| 15 | 1 | -6 | 46 | -26 | L | Orbital Gyrus | 11 |
| 11 | 3 | 60 | 16 | 6 | R | Inferior Frontal Gyrus | 45 |
| 11 | 4 | -68 | -6 | 8 | L | Superior Temporal Gyrus | 22 |
| 11 | 6 | -4 | 64 | -8 | L | Superior Frontal Gyrus | 10 |
| 10 | 1 | -2 | 40 | -20 | L | Orbital Gyrus | 11 |
| 9 | 7 | 40 | -12 | -30 | R | Inferior Temporal Gyrus | 20 |
| 8 | 6 | -3 | 63 | -5 | L | Medial Frontal Gyrus | 10 |
| 8 | 8 | -54 | 10 | 26 | L | Inferior Frontal Gyrus | 9 |
| 7 | 7 | 37 | -10 | -30 | R | Uncus | 20 |
| 7 | 8 | -53 | 8 | 26 | L | Inferior Frontal Gyrus | 9 |
| 4 | 1 | -4 | 30 | -12 | L | Medial Frontal Gyrus | 11 |
| 4 | 2 | 52 | -11 | 21 | R | Postcentral Gyrus | 43 |
| 4 | 3 | 56 | -5 | -4 | R | Middle Temporal Gyrus | 21 |
| 4 | 4 | -61 | -28 | 21 | L | Postcentral Gyrus | 40 |
| 4 | 5 | 24 | -64 | 24 | R | Precuneus | 31 |
| 4 | 5 | 7 | -67 | 21 | R | Precuneus | 31 |
| 4 | 6 | -15 | 61 | -10 | L | Superior Frontal Gyrus | 11 |
| 4 | 7 | 31 | -6 | -32 | R | Uncus | 20 |
| 4 | 8 | -55 | 12 | 22 | L | Inferior Frontal Gyrus | 45 |

**Table S5.** Clusters of specificity in Alzheimer's disease (AD).
BF Value = Bayes' Factor Value

| *BF Value* | *Cluster* | *x* | *y* | *z* | *Side* | *Label* | *BA* |
|---|---|---|---|---|---|---|---|
| 21 | 1 | 30 | -58 | 10 | R | Parahippocampus | 30 |
| 13 | 5 | -64 | -46 | 40 | L | Inferior Parietal Lobule | 40 |
| 12 | 2 | 34 | -36 | 0 | R | Caudate (Caudate Tail) | |
| 10 | 8 | -48 | -62 | 52 | L | Inferior Parietal Lobule | 0 |
| 9 | 2 | 29 | -37 | 0 | R | Hippocampus | |
| 8 | 7 | 62 | -38 | 40 | R | Inferior Parietal Lobule | 40 |
| 7 | 7 | 53 | -50 | 34 | R | Supramarginal Gyrus | 40 |
| 6 | 3 | 24 | -8 | -10 | R | Amygdala | |
| 6 | 4 | -26 | -10 | -10 | L | Amygdala | |
| 6 | 6 | -32 | -36 | -2 | L | Hippocampus | |
| 6 | 6 | -32 | -34 | -4 | L | Hippocampus | |
| 5 | 1 | 18 | -60 | 20 | R | Posterior Cingulate | 31 |
| 5 | 3 | 19 | -9 | -10 | R | Amygdala | |
| 5 | 4 | -24 | -7 | -16 | L | Amygdala | |
| 5 | 8 | -41 | -53 | 55 | L | Inferior Parietal Lobule | 40 |
| 4 | 1 | 23 | -61 | 11 | R | Posterior Cingulate | 30 |
| 4 | 2 | 28 | -50 | -5 | R | Parahippocampus | 19 |
| 4 | 3 | 21 | -20 | -16 | R | Parahippocampus | 35 |
| 4 | 4 | -17 | -11 | -10 | L | Parahippocampus | 28 |
| 4 | 5 | -57 | -54 | 29 | L | Superior Temporal Gyrus | 39 |
| 4 | 6 | -26 | -42 | -5 | L | Parahippocampus | 36 |
| 4 | 7 | 53 | -46 | 30 | R | Supramarginal Gyrus | 40 |
| 4 | 8 | -34 | -50 | 50 | L | Superior Parietal Lobe | 7 |

**Table S6.** Clusters of Pain specificity.
BF Value = Bayes' Factor Value

| *BF Value* | *Cluster* | *x* | *y* | *z* | *Side* | *Label* | *BA* |
|---|---|---|---|---|---|---|---|
| 11 | 2 | -36 | -20 | 14 | L | Insula | 13 |
| 10 | 1 | 36 | -18 | 18 | R | Insula | 13 |
| 9 | 3 | 2 | 4 | 36 | R | Cingulate Gyrus | 24 |
| 9 | 3 | -2 | 4 | 37 | L | Cingulate Gyrus | 24 |
| 7 | 5 | 8 | -18 | 12 | R | Thalamus (Medial Dorsal Nucleus) | |
| 7 | 5 | 9 | -16 | 12 | R | Thalamus (Medial Dorsal Nucleus) | |
| 7 | 6 | -12 | -14 | 8 | L | Thalamus (Medial Dorsal Nucleus) | |
| 7 | 6 | -13 | -14 | 7 | L | Thalamus (Ventral Lateral Nucleus) | |
| 6 | 7 | 48 | 46 | -10 | R | Middle Frontal Gyrus | 47 |
| 5 | 8 | 30 | -46 | -39 | R | Cerebellar Tonsil (Cerebellum) | |
| 4 | 1 | 27 | -12 | 21 | R | Claustrum | |
| 4 | 1 | 40 | -7 | 14 | R | Insula | 13 |
| 4 | 2 | -27 | 0 | 10 | L | Putamen | |
| 4 | 2 | -44 | -12 | 14 | L | Insula | 13 |
| 4 | 3 | -17 | -6 | 38 | L | Cingulate Gyrus | 24 |
| 4 | 4 | -27 | -56 | -26 | L | Culmen (Cerebellum) | |
| 4 | 5 | 10 | -12 | 0 | R | Thalamus | |
| 4 | 6 | -7 | -11 | -1 | L | Thalamus | |
| 4 | 7 | 39 | 43 | -6 | R | Middle Frontal Gyrus | 11 |
| 4 | 7 | 40 | 44 | -1 | R | Inferior Frontal Gyrus | 10 |
